# Identification and physiological significance of temporal NFκB signaling codewords deployed by macrophages to classify immune threats

**DOI:** 10.1101/2020.05.23.112862

**Authors:** Brooks Taylor, Adewunmi Adelaja, Yi Liu, Stefanie Luecke, Alexander Hoffmann

## Abstract

Acute and chronic inflammatory pathologies involve misregulation of macrophage functions. Physiologically, macrophages are immune sentinels that initiate inflammatory responses via the transcription factor NFκB. The temporal pattern of NFκB activity determines which genes are expressed, suggesting that a temporal signaling code specifies a stimulus-appropriate immune response. To identify the signaling codewords, we developed tools to enable high-throughput analysis of live, primary macrophages responding to host- and pathogen-derived stimuli. An information-theoretic workflow identified six dynamical features that constitute codewords that convey stimulus information to the nucleus. In particular, “oscillatory” trajectories are a hallmark of the responses to host cytokine TNF. Remarkably, examining macrophages derived from a systemic autoimmune disease model suggests that confusion of two NFκB signaling codewords, and thus miscoding of TNF as a pathogen-derived stimulus, may underlie sporadic inflammatory pathology. Overall, this study identifies six codewords of the temporal NFκB signaling code for classifying immune threats and demonstrates their biological significance.

## INTRODUCTION

Autoimmune pathologies are characterized by the presence of auto-antibodies and immune attack of specific tissues, but the etiology is not uniform ^1^. One cause may be found in errors in the negative selection of auto-reactive B-cell or T-cell clones in secondary lymphoid organs ^1^; another contributor may be inappropriate immune activation by immune sentinel cells ^1^. Sjögren’s syndrome (SS) is a systemic autoimmune disorder, which is characterized by progressive destruction of tissues exposed to the environment, such as eye, mouth and throat, and skin rashes ^1^. Interestingly, several genetic variants in regulators of the inflammatory transcription factor NFκB are associated with SS patients ^2–5^ and a mouse strain containing similar variants recapitulates some of the SS pathognomonic characteristics ^6^. However, it remains unknown how these alleles affect control of NFκB dynamics.

Macrophages are immune sentinel cells that respond to pathogen invasion and tissue injury by initiating and coordinating both local and system-wide immunity ^7^. These cells are ubiquitously distributed in tissues ^8^ and can sensitively detect inflammatory cytokines and pathogen-associated molecular patterns (PAMPs), which indicate viral, bacterial, or fungal invasion ^9^. Immune activation must be appropriate to each stimulus: the functional response to the cytokine TNF must be distinct from the response to a pathogen; further, the needs of a macrophage responding to bacterial or viral invasion are distinct.

The temporal coding hypothesis posits that information about the extracellular stimulus is represented in the time-domain, i.e. the temporal pattern of a signaling activity ^10–12^. Biochemical studies in primary fibroblasts showed that the temporal pattern of NFκB RelA activity is stimulus-specific at the cell population level ^13,14^, and that it controls the expression of immune response genes ^18, 15^. Although pioneering single-cell microscopy studies confirmed complex temporal patterns ^16–18^, they relied upon fluorescent-protein-NFκB RelA fusion proteins ectopically expressed in immortalized cell lines (Supplementary Table 1). Potential artefacts arising from ectopic expression of a reporter-effector protein have been reported ^19,20^, and prolonged cell culture adaptation of immortalized cell lines was found to diminish their responsiveness to immune threats ^21^. These limitations have not allowed the literature to explore the biological significance of temporal coding in primary immune cells and whether it is a useful concept for understanding immune pathology. Reasons for why no studies of single-cell NFκB trajectories in primary macrophages have been reported thus far include challenges associated with imaging knockin fluorescent protein reporters that are not overexpressed and reliable high-throughput image analysis of morphologically heterogeneous cells.

Here, we report a new mVenus-RelA knockin mouse strain and a high-throughput imaging and analysis workflow to investigate the NFκB temporal code in single, primary macrophages. An information-theoretic approach was used to identify dynamical features of the NFκB trajectories that convey information about the extra-cellular stimulus to the nucleus. Teaching these codewords to a machine demonstrated their sufficiency and requirement for ligand and dose identification. Indeed, examination of a SS mouse model revealed confusion of specific signaling codewords and suggested that such confusion may contribute to the etiology of systemic autoimmune diseases. Finally, mathematical modeling allowed us to identify the molecular circuit design principles that enable encoding of these newly identified signaling codewords, and confirmed that ‘oscillations’ are a hallmark of responses to the host cytokine TNF, in contrast to PAMPs transduced by the signaling adaptor MyD88.

## RESULTS

### Primary macrophages show immune threat ligand- and dose-specific NFκB dynamics

To study temporal patterns of nuclear NFκB in primary macrophages in response to prototypical immune threats (Fig. 1A) at single-cell resolution, we generated a mouse strain expressing a mVenus-RelA fusion protein (Supplementary Fig. 1A, B), similar to a previous GFP-RelA design ^22^ whose low fluorescence limited experimental studies ^23^. Macrophages, differentiated from primary bone-marrow cells derived from homozygous mVenus-RelA mice, showed normal levels of nuclear NFκB binding activity (Supplementary Fig. 1C). Upon stimulation with a variety of different ligands and doses, and time-lapse imaging over 21 hours (Fig. 1B), the amount of nuclear NFκB fluorescence was quantitated in single cells using a fully automated image processing pipeline that enabled tracking of live cells using minimal levels of a nuclear marker ^24,25^ and label-free identification and segmentation of cell cytoplasm. The live-cell imaging and image processing proved robustly reproducible in biological replicates (Supplementary Fig. 1D), and independent of image frame location (Supplementary Fig. 1E). We noted striking differences in the NFκB dynamics induced by prototypical PAMP (LPS) and cytokine (TNF) stimuli (Movie S1), apparent at the single-cell level (Fig. 1C). TNF induced oscillatory translocations between cytoplasm and nucleus that rapidly became desynchronized, matching biochemical data ^15^. By contrast, LPS induced more than 4 hours of sustained nuclear localization that also matched biochemical data from primary fibroblasts ^13,14^.

**Figure 1.**
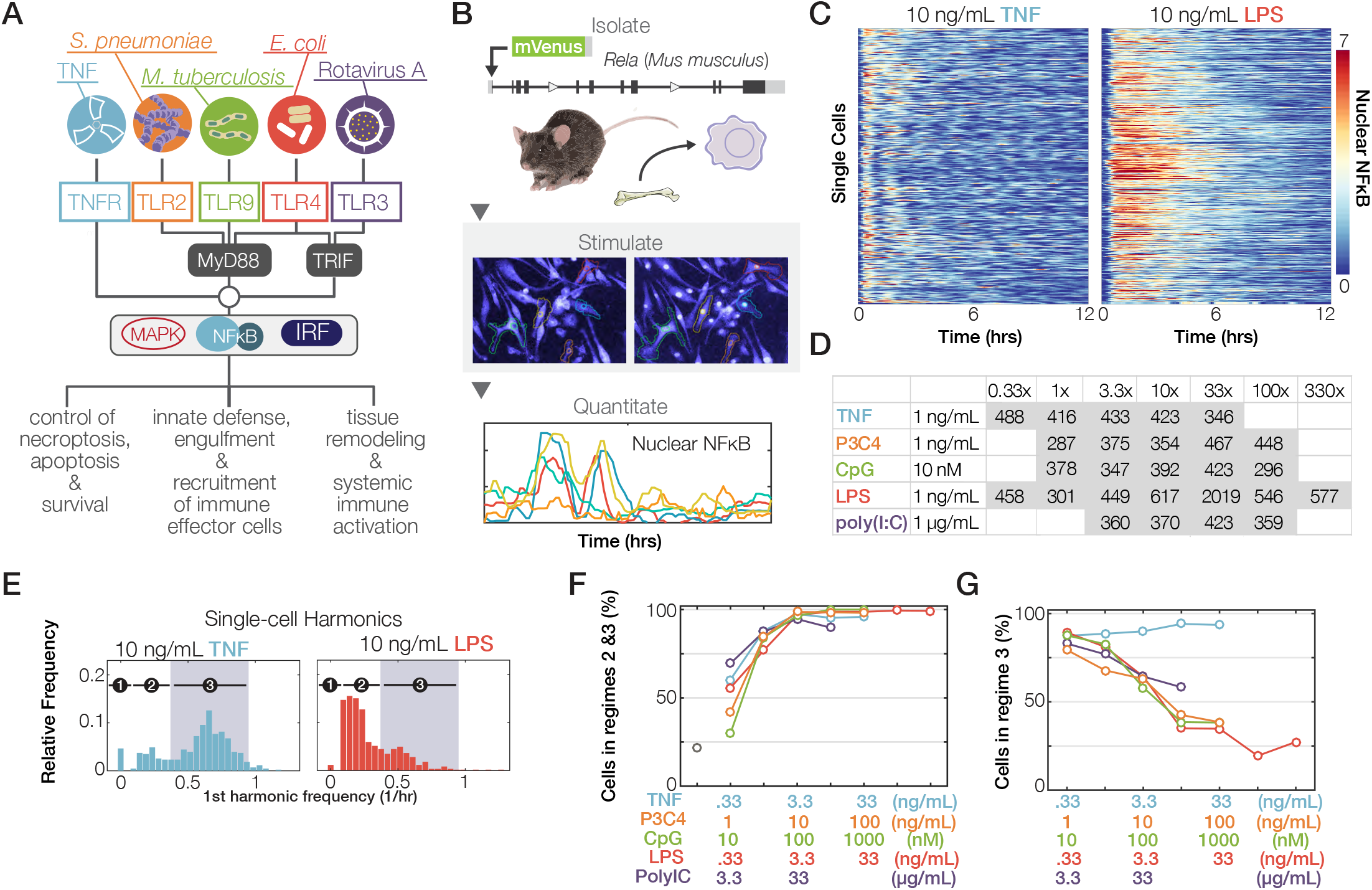
Complex NFκB dynamics induced by diverse immune threats. (A) Schematic of the innate immune signaling network activating NFκB. Environmental information is transmitted via ligand-specific signaling pathways that converge on a few key transcription factors, including NFκB, but produce stimulus-specific physiological responses. (B) Workflow diagram: a knockin mouse line expressing mVenus-RelA was generated. Bone marrow-derived macrophages (BMDMs) were differentiated, imaged, tracked, and quantified in multiple stimulus conditions. (C) Single-cell heatmaps of fluorescent nuclear NFκB levels over time, in BMDMs expressing endogenously tagged mVenus-RelA, in response to 10 ng/mL TNF or LPS. Each row is one cell’s trajectory. (D) Table indicating the number of single cell trajectories quantified in each indicated experimental condition. This analysis involved 12,203 cell trajectories produced by quantifying more than 3 million cell images. More details in Supplementary Table 2. All single-cell imaging data was confirmed, here and elsewhere, with at least two independent experiments per condition. (E) First-harmonic distributions for other stimuli. Shaded region corresponds to the period of 1-2.2 hrs that is characteristic of NFκB oscillations. (F) Fraction of cells in which a response is detected, by stimulus and dose. (G) Fraction of responder cells that show characteristic NFκB oscillations.

With an experimental workflow established, we recorded NFκB translocation dynamics in response to a large number of stimulation conditions, encompassing TNF and four different PAMPs, associated with diverse bacterial and viral pathogen classes (the TLR ligands CpG (TLR9), Pam3CSK4 (TLR1/2, referred to as P3C4), LPS (TLR4), and Poly(I:C) (TLR3)) each tested at 4-7 concentrations covering a 10^2^ to 10^3^-fold range. In each condition, 300-600 cells were examined with at least two preparations of BMDMs, thus constituting a total dataset of 12203 single-cell trajectories captured with more than 3 million cell images and associated NFκB activity datapoints (Fig. 1D, Supplementary Table 2).

Given the NFκB trajectory, each cell was classified based on its first harmonic frequency profile generated by Fourier analysis (Fig. 1E) as either unresponsive (regime 1), responsive but non-oscillatory (regime 2), or oscillatory (regime 3) with a period of 1.1-2.2 hours characteristic of NFκB oscillations ^26^. Analysis of the data indicated that the lowest stimulus concentration activated about half the cells but that a log_10_ increase activated almost all (Fig. 1F). Plotting the percentage of cells classified as oscillators in responders, we found that the host factor TNF elicited oscillatory dynamics regardless of dose (Fig. 1G). While the number of peaks increased with increasing doses of TNF, the period remained constant (Supplementary Fig. 1F). In contrast, PAMPs produced largely non-oscillatory responses at high ligand concentrations (Fig. 1G). Unlike experimental systems with ectopically expressed RelA, which produced a first peak of NFκB activity that is much higher than later peaks ^17,18,27^, the response of primary macrophages to TNF showed a constant, gradual fall-off in amplitude (Supplementary Fig. 1G).

### Informative dynamical features are identifiable

Oscillations are just one dynamical feature by which complex time course trajectories can be characterized. We developed a method for identifying dynamical features that are associated with stimulus- and dose-specific NFκB trajectories. We constructed a novel multivariate information-theoretic algorithm, based on an estimate of channel capacity ^24,28^. In addition to the primary timeseries data, we considered 918 derived metrics (Supplementary Table 3) such as integrals, derivatives, peak activities, durations, or frequencies (Fig. 2A). Our algorithm searched this library for combinations of metric that maximized channel capacity (Supplementary Fig. 2A), iteratively expanding the number of metrics within each combination from two up to ten.

**Figure 2.**
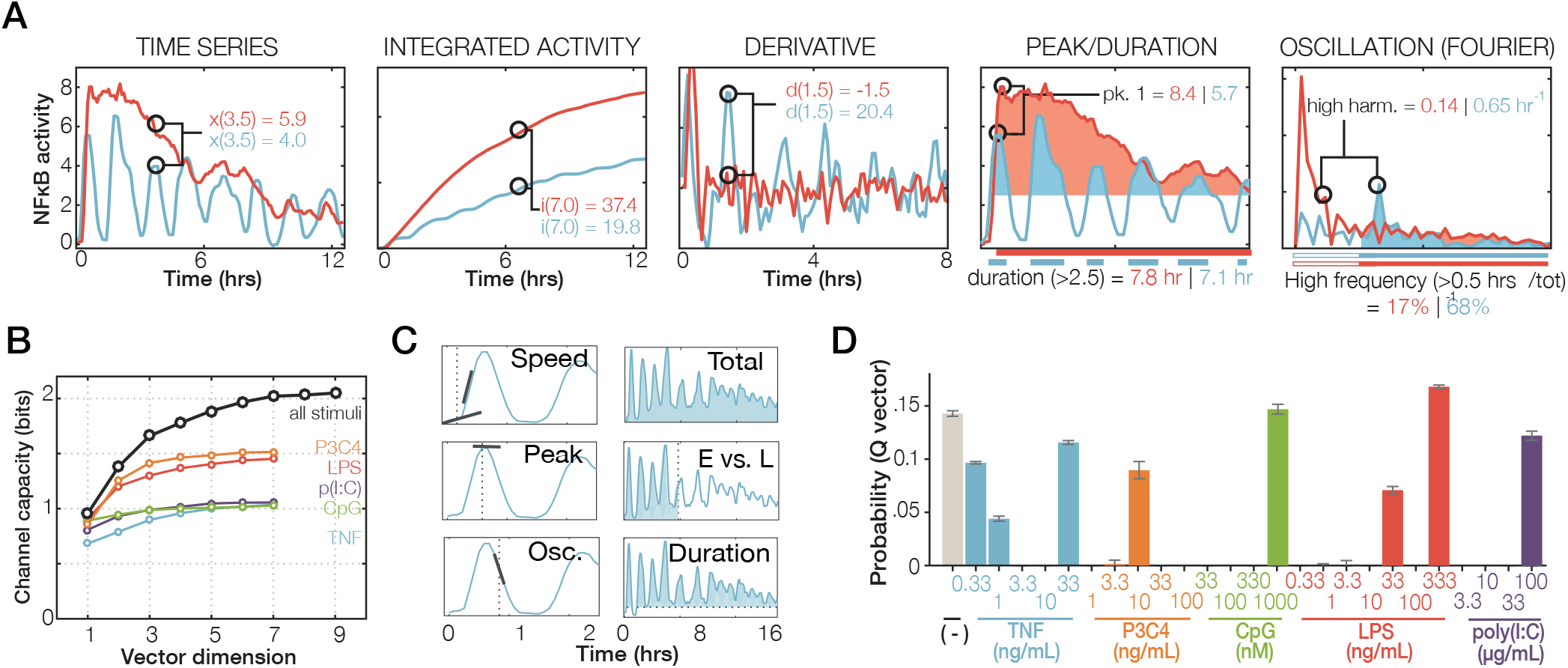
Informative features within complex NFκB dynamics. (A) Examples of metrics to be employed in an information theoretic analysis. Two single-cell NFκB responses (to LPS in red, and to TNF in blue) are shown. All NFκB trajectories were characterized using 918 metrics (Supplementary Table 3). (B) Channel capacity as a function of the number of most informative metrics (Supplementary Table 4), either using the entire dataset of all ligand types and doses (black line) or using the dose response data for each indicated ligand. (C) ynamical features that are informative about ligand and dose, as revealed by the metrics selected by the information theoretic analysis. E: early activity; L: late activity. (D) Average probability distribution from the channel capacity calculations using all optimal vectors. Probabilities sum to 1 and indicate the input distribution that leads to a computationally-maximized mutual information.

First, we considered the available dose response dataset for each ligand separately. Combinations of five metrics were sufficient to capture the mutual information of dose responses, with TNF, CpG and Poly(I:C) achieving about 1 bit and LPS and Pam3CSK4 about 1.5 bits (Fig. 2B), in agreement with previous reports for TNF and LPS ^24,28^. When considering all ligands tested (26 dose-ligand conditions), the calculated channel capacity was markedly higher (>2 bits) and required a seven-dimensional vector to yield ≥ 95% of the maximum measured information content (Supplementary Table 4).

Of these most informative metrics identified across the full dataset (Fig. 2C), two defined the activation speed (1), one defined the peak amplitude (2), another defined the post-induction repression, a distinguishing feature of oscillatory versus non-oscillatory trajectories (3), one defined the accumulated activity (integral) at a late time (4), one was measure of the degree to which NFκB activity is ‘front-loaded’ (5); and one defined the total duration of NFκB activity above a low threshold (6). Thus the information-theoretic analysis identifies six NFκB dynamical features that are informative about the stimulus ligand and dose. Plotting three features allowed for only incomplete separation of ligands (Supplementary Fig. 2B).

Further analysis of the channel capacity calculations indicated that the highest dose generally provided the most ligand-specific information (Fig. 2D). Indeed, when we restricted the calculation to only the highest dose of each of our five ligands we still obtained a channel capacity of 1.86 bits. It is remarkable that unlike the dose response profiles of pharmacological agents, which tend to show cross-reactivity at high doses, ligand-specific signaling dynamics occur at highest doses, indicating that there are true differences in the signal processing characteristics of receptor-associated signaling pathways.

### Machine learning of NFκB codewords distinguishes stimuli

The six NFκB dynamical features, identified as conveying information about the extra-cellular stimulus to the nucleus, represent potential codewords of the temporal NFκB signaling code. Visualizing codeword deployment for the five ligands at high doses (Fig. 3A, Supplementary Table 5), the speed of activation is generally high for Pam3CSK4 and LPS-triggered signaling, but low for CpG and Poly(I:C), and intermediate for TNF; peak amplitude is high for Pam3CSK4, CpG and LPS and lower for TNF and Poly(I:C); the oscillatory content is highest for TNF compared to any of the PAMPs; the amount of total activity is highest for LPS followed by Poly(I:C) and Pam3CSK4, but lower for TNF and CpG; the total duration, in contrast, is high for TNF and Poly(I:C), and relatively low for Pam3CSK4, CpG, and LPS; and the fraction of the activity that is early is much higher for TNF, Pam3CSK4, and LPS than Poly(I:C), with CpG being intermediate. Similarly, we find that different doses of the same ligand may deploy the codewords differentially (Supplementary Fig. 3A). For example, while the peak activity is generally positively correlated with dose ^29^, the duration of activity increases with increasing doses of TNF or LPS, but decreases with increasing doses of CpG.

**Figure 3.**
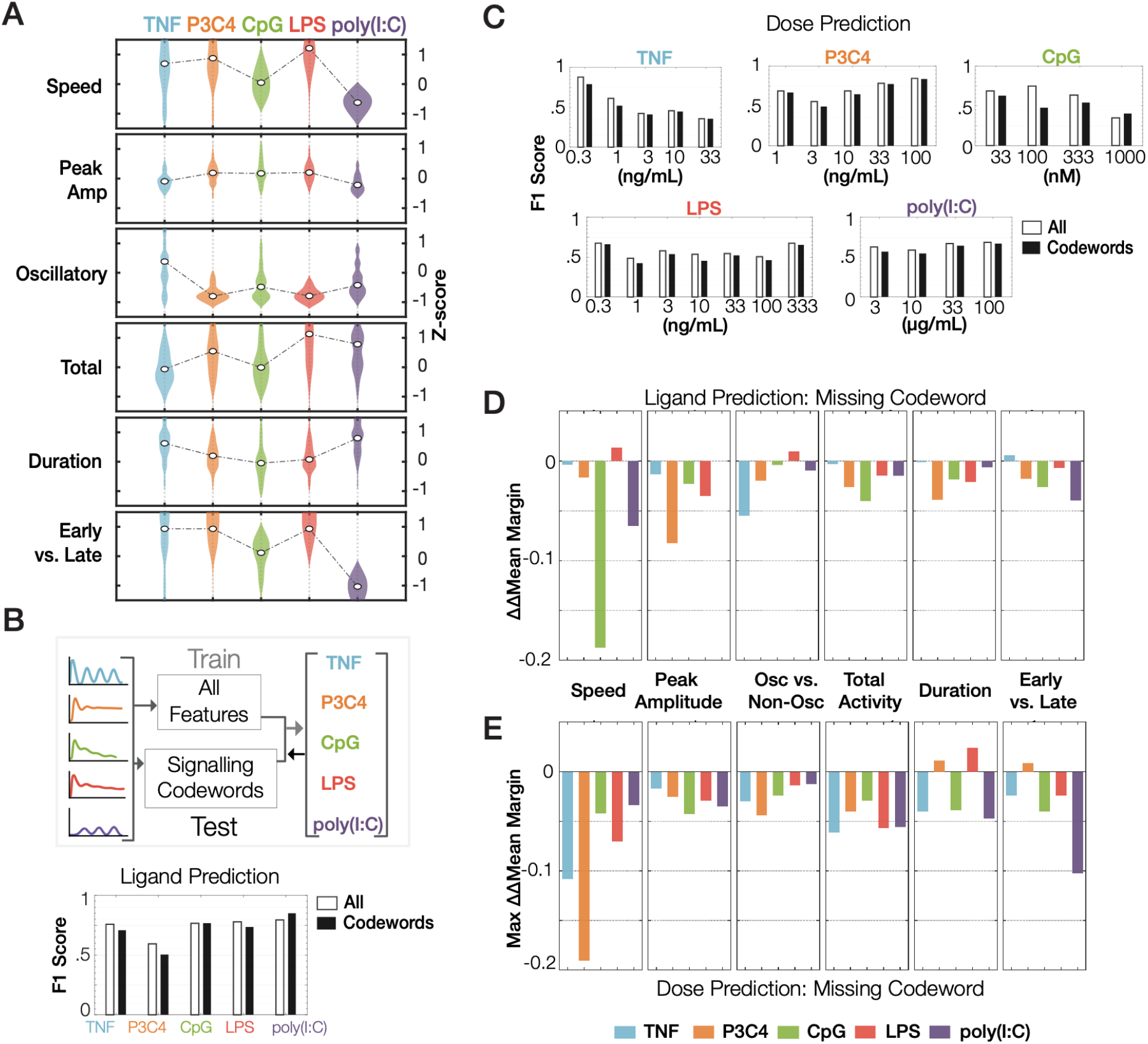
Six NFκB Signaling ‘codewords’ are sufficient to classify immune threats. (A) Violin plots of dynamical features that optimally encode stimulus-specific NFκB dynamics: activation speed, peak amplitude, oscillatory dynamics, total activity, duration, and ratio of early to late activity. These are termed “signaling codewords”, and they are deployed in a stimulus-specific manner as shown. (B) Top: Schematic of supervised machine learning approach to predict ligand identity using NFκB dynamics. Bottom: F1 scores (harmonic mean of precision and recall) of ligand predictions using either all features or signaling codewords alone. (C) F1 score of dose predictions for each indicated ligand using either all features or only codewords. (D) The effect of each signaling codeword on the certainty of ligand prediction: The loss in classification confidence when the indicated codeword is missing from the set of six (versus all features). Mean classification margin: the difference between the true ligand probability and maximum false ligand probability, ΔMean Margin: difference in mean classification margin of codon classifier vs. all predictors classifier; ΔΔMean Margin: difference in ΔMean Margins when using codon classifier with all six codewords or with classifiers with indicated codewords missing vs. all predictor classifiers. (E) The effect of each signaling codeword on the certainty of dose prediction for each ligand: The loss in classification confidence when the indicated codeword is missing from the set of six (versus all features).

To determine whether NFκB signaling codewords suffice to distinguish these ligands, we used supervised machine learning and trained an ensemble-of-decision-trees model either with all 918 metrics or the set of six signaling codewords (Supplementary Fig. 3B). We chose this classification algorithm because of its performance and interpretability ^30,31^. Assessing prediction performance, we found that F1 scores (harmonic mean of precision and recall, a measure of specificity and sensitivity of the predictions) were remarkably similar for predictions generated using all metrics or just signaling codewords (Fig. 3B, Supplementary Table 6A). Other performance measures confirm this conclusion (Supplementary Fig. 3C), indicating that signaling codewords alone suffice to distinguish NFκB ligands. Using the same approach, we examined whether signaling codewords suffice to distinguish the doses of each ligand. The differences in F1 scores of dose predictions generated by classifiers trained using all features versus six signaling codewords were minimal (Fig. 3C, Supplementary Table 6B).

We quantified the certainty of stimulus classification (classification margin; difference of the true ligand probability and maximum false ligand probability) using all features, signaling codewords, and subsets of signaling codewords (Supplementary Fig. 3D). To examine the necessity of each signaling codeword, we computed ΔΔMean Margin, which is the difference in the mean classification margin (using either all features or merely the six signaling codewords, ΔMean Margin) subtracted by the difference in the mean classification margin when one codeword is removed. This analysis revealed the stimulus-specific dependence of classification certainty on signaling codeword: speed is important in classifying CpG and Poly(I:C), peak amplitude is important for classifying Pam3CSK4, and oscillatory dynamics are important for classifying TNF (Fig. 3D).

To examine the necessity of each signaling codeword in distinguishing doses, we quantified the ΔΔMean Margin across all doses for each ligand (Fig. 3E, Supplementary Fig. 3E). The maximum ΔΔMean Margin across all doses of each ligand revealed that speed is important to distinguish doses of TNF, Pam3CSK4, and LPS, and early-vs.-late activity is important to distinguish doses of Poly(I:C). Furthermore, this analysis suggests that the importance of a signaling codeword for classifying a ligand may differ from its importance in distinguishing the doses of that ligand (Fig. 3D and E). Using binary classification of stimulated condition vs. vehicle control indicated that ligand identification increases with the dose of the stimulus (Supplementary Fig. 3F), confirming the results of the information theoretic analysis (Fig. 2D).

### Increased codeword confusion in autoimmune disease

The availability of a validated machine learning classifier allowed us to quantify not only how precise stimulus identification is, but which other stimuli a given stimulus may be confused with. We characterized the points of confusion by quantifying classification accuracy (precision) in the matrix of five ligands, choosing their highest doses as they are most distinguishable (Fig. 4A, Supplementary Fig. 4A). Correct classification of ligand identities occurred in the majority, but misclassifications (off-diagonal values) were not uniformly distributed. For example, while confusion of viral PAMP Poly(I:C) and bacterial PAMP LPS was rare, it was more common between the bacterial PAMPs, LPS and Pam3CSK4. Indeed, when we grouped ligands into their source classes such as host (cytokine), bacteria, or virus, we found that bacteria-derived ligands are reliably distinguished and show little confusion with either virus- or host-derived ligands (Fig. 4B, Supplementary Fig. 4B).

**Figure 4.**
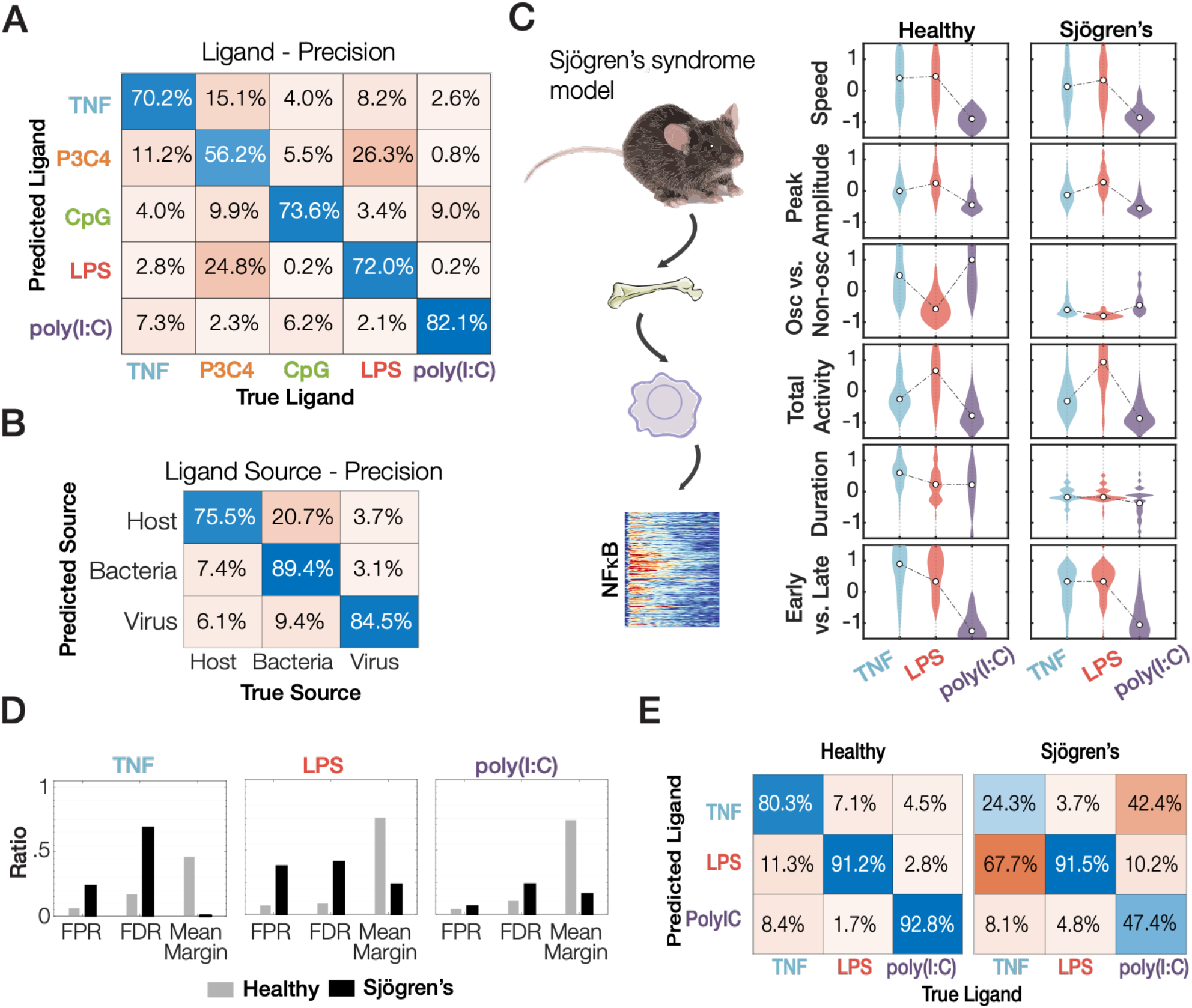
A Sjögren’s syndrome mouse model shows more confusion in classifying immune cytokine TNF and immune threat LPS based on NFκB dynamics. (A) Confusion matrices showing classification precision of ligand identity information. The machine learning model correctly identifies the ligand identity given an NFκB trajectory a majority of the time with the primary confusion being between bacterial ligands Pam3CSK4 and LPS most apparent. (B) Confusion matrices showing classification precision of ligand source information. Bacterial ligands are generally correctly identified as such. (C) Testing ligand confusion in macrophages isolated from a Sjögren’s disease model mouse ^32^. Violin plots depicting the signaling codewords deployed by macrophages, derived from healthy or Sjögren’s mice, stimulated with TNF, LPS or Poly(I:C). (D) Classification of ligand identity in healthy and Sjögren mouse model macrophages by a machine learning classifier trained on healthy macrophage data: False positive rate (FPR), false discovery rate (FDR), and mean margin. (E) Confusion matrices for sensitivity of recall for the healthy and Sjögren’s macrophage data.

We assessed whether a mouse model of Sjögren’s syndrome (SS) ^32^, which harbors genetic variants in the regulatory region of the NFκB regulator IκBα, may be associated with signaling codeword confusion, such that cells exposed to one stimulus might in fact miscommunicate the presence of a different stimulus to nuclear target genes. We bred our mVenus-RelA reporter into this mouse model and then derived bone marrow-derived macrophages for stimulation with the cytokine TNF, the bacterial PAMP LPS, and the viral PAMP poly(I:C). Unlike macrophages from healthy mice, these SS macrophages showed non-oscillatory NFκB trajectories in response to all stimuli (Supplementary Fig. 4C - E). Visualizing the six codewords revealed that the stimulus-specific deployment of particular NFκB signaling codewords was impaired in macrophages from the Sjögren’s mouse model (Fig. 4C). The stimulus-specificity of the “oscillatory” codeword was markedly diminished in SS macrophages, and the stimulus-specificity of the “duration” and the “early vs. late” codewords was also affected.

Then, we examined the accuracy of stimulus classification using the ensemble-of-decision trees algorithm (Supplementary Table S6C). The mean margin scores of ligand classification were greatly diminished in SS macrophages, concomitant with an elevation in the false positive and false discovery rates for TNF and LPS (Fig. 4D). Furthermore, the sensitivity of TNF and poly(I:C) classification in SS macrophages was greatly diminished (24.3%/47.4%, respectively, versus 80.3%/92.8% in healthy controls, Fig. 4E), as there is increased confusion between Poly(I:C) versus LPS, and TNF versus LPS. These analyses indicate that SS macrophages have diminished ability to generate stimulus-specific NFκB signaling dynamics and suggest that signaling codeword confusion and mistranslation may play a role in the etiology of sporadic inflammatory diseases.

### Molecular circuits that produce signaling codewords

Having identified essential dynamical features of NFκB activity for encoding ligand identity and dose, we sought to understand the molecular mechanisms that provide for the diversity of stimulus-specific dynamics. The known topology of the NFκB network is that signals emanating from receptor-associated signaling modules converge to activate canonical IKK, which functions as the input to the IκB-NFκB signaling module whose most prominent regulator is IκBα (Fig. 5A, Supplementary Fig. 5A). A prominent signaling codeword that distinguishes the cytokine TNF from PAMPs is the oscillatory content. Using macrophages from an IκBα-deficient mouse (interbred with the mVenus-RelA reporter, see methods), we found at the single-cell level that oscillatory dynamics are dependent on IκBα negative feedback (Fig. 5B), in agreement with prior population level experiments ^15,33^.

**Figure 5.**
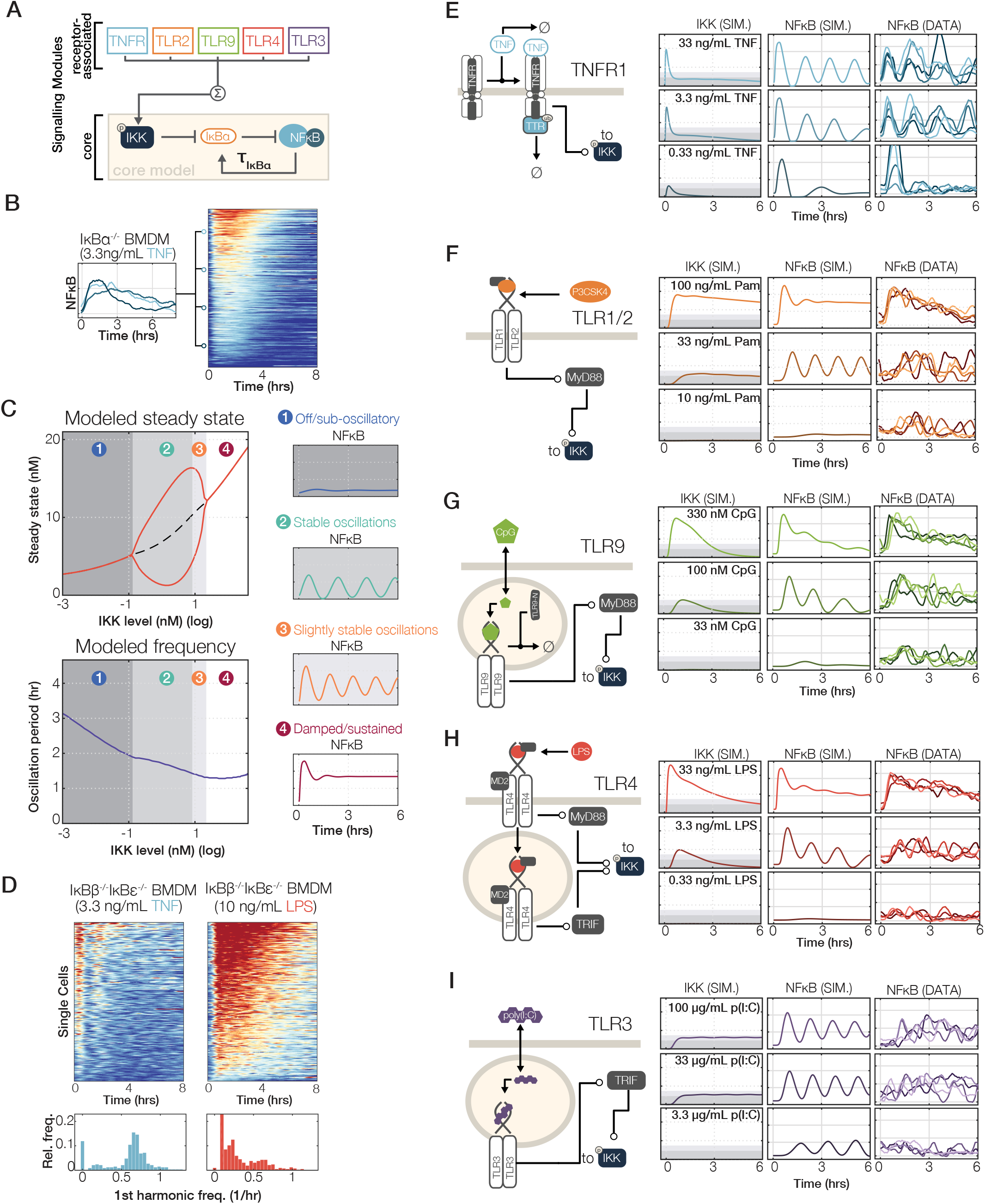
Kinetic models of receptor-associated signaling modules share circuit design principles that generate NFκB signaling codewords in a stimulus-specific manner. (A) A simple schematic suggesting that NFκB control is mediated by two regulatory networks: The core IκBα-NFκB signaling module is downstream of receptor-associated signaling modules. Receptor-associated signaling modules determine IKK activity over time. Within the core module, IKK activity destabilizes IκBα, freeing NFκB to translocate to the nucleus, where it induces expression of IκBα. (B) The IκBα-feedback is required for generating the oscillatory component of NFκB dynamics characteristic of the response to TNF. Single-cell trajectories and heatmaps of NFκB responses to 3.3 ng/mL TNF in BMDMs derived from a mVenus-RelA, IκBα-deficient mouse. (C) A computational model predicts bifurcating behavior in NFκB dynamics based on the level of IKK activation. Left: model steady state values and primary oscillation frequency are shown as a function of sustained IKK level. Right: single simulated trajectories of IKK and NFκB activation, at each of four regimes identified in the steady state diagram. (D) The IκBα-feedback is sufficient sustain non-oscillatory NFκB characteristic of the response to LPS. Single cell heatmaps of NFκB responses to 3.3 ng/mL TNF and 10 ng/mL LPS in BMDMs derived from a mVenus-RelA, *IκBβ^−/−^IκBε^−/−^* mouse. Below each heatmap is a histogram of each cell’s first harmonic showing relative proportions of oscillatory cells. (n>400 individual cells for each experiment, representative of two independent replicates). (E) - (I) Simplified schematics showing salient features of TNF, TLR1/2, TLR9, TLR4, and TLR3 signaling pathways, and the simulated IKK and NFκB activity (left/middle), and four measured median cell NFκB trajectories (right) at each of three log-spaced (TNF and TLR4) or four half-log spaced (TLR9, TLR1/2, and TLR3) doses of each receptor’s cognate ligand. The complete reaction sets of the model are described in the Methods and Supplementary Table 7.

Then, we examined whether the IκBα feedback loop may also mediate non-oscillatory responses characteristic of PAMPs or whether other IκB isoforms may be required. After adapting the mathematical model of the negative-feedback containing IKK-IκBα-NFκB signaling module to the primary macrophage (see Methods), we examined its dynamical properties using Hopf-bifurcation analysis, specifically the propensity for oscillatory responses as a function of the magnitude of IKK activity (Fig. 5C). The first bifurcation point defines the threshold between (1) “off” (indistinguishable from baseline activity) and (2) an oscillatory steady-state. As IKK activity increases, oscillation troughs rise in amplitude (3) though the period changes little. The second bifurcation point occurs as the system shifts to highly damped oscillations (4). Our analysis thus led to the prediction that non-oscillatory NFκB responses of LPS are not mediated by other IκB isoforms (IκBβ and IκBε), as previously hypothesized ^34,35^, but that the NFκB-IκBα feedback circuit alone could sustain such non-oscillatory behavior. To test this hypothesis, we bred our RelA-mVenus reporter into *IκBβ^−/−^IκBε^−/−^* mice and gathered single-cell responses to TNF and LPS (Fig. 5D). In this genotype, TNF induced an even higher fraction of oscillatory cells (95% versus 75% in wild-type, Fig. 1G), while LPS responses were, as before, largely non-oscillatory. We conclude that both oscillatory and non-oscillatory NFκB dynamics may be generated by the IκBα-NFκB signaling module; the deployment of the “oscillatory” signaling codeword is determined merely by controlling the amount of IKK activity over time.

To build a full, multi-stimulus model, capable of generating proper IKK activity timecourses in response to any of the ligands and doses used in this study, we carefully examined the regulatory mechanisms associated with each ligand receptor (Supplementary Fig. 5A), and drafted ordinary differential equations to describe them. Parameter values were based on prior literature (Supplementary Table 7) and adjusted to produce model simulations of NFκB that qualitatively matched trajectories of median-responding cells in each tested condition (Fig. 5E-I). For TNF and LPS, available literature datasets on receptor and IKK dynamics were fit (Supplementary Fig. 5B, C). Within the core IKK-IκB-NFκB module, multi-parameter sampling confirmed that the oscillatory-non-oscillatory distinction based on the magnitude of IKK activity was a robust feature (Supplementary Fig. 5D). In addition, model-simulated IKK trajectories (Fig. 5E-I) were tested at key timepoints using immunoblotting of the active, phosphorylated IKK species (Supplementary Fig. 6A - E) and revealed a general concordance in this semi-quantitative comparison. While this increases our confidence in the insights derived from the model, we cannot rule out alternative models or mechanisms.

Signaling within each signaling module is governed largely by the kinetic properties of a few constituents such as ligand half-life, receptor downregulation and replenishment, and the dose response properties of the receptor-associated signaling adaptor. For example, in the case of TNF, rapid receptor downregulation and short ligand half-life ^36,37^ diminish IKK activity into a regime that allows for deployment of the ‘oscillatory’ codeword and the dose-dependent deployment of the ‘duration’ codeword, respectively (Fig. 5E). For Pam3CSK4 and CpG (Fig. 5F and G), the signaling characteristics of cooperative adaptor interactions lead to digital dose response behavior ^21^ and low values for the ‘oscillatory’ (due to high IKK activity) and ‘duration’ codewords at high doses. In the case of LPS-TLR4 (Fig. 5H) the combination of ultrasensitive and linear dose response behavior of MyD88 and TRIF adaptors ^21,38^, aided by CD14-mediated TLR4 internalization ^39^, provide for dose-dependent deployment of the codewords of ‘oscillatory’ and ‘total activity’. In contrast, endosomal availability of TLR3 and Poly(I:C) ^40^ limit the ‘response speed’ codeword but allow for long ‘duration’ (Fig. 5I). Overall, the comparison of five signaling modules revealed shared molecular circuit design principles whose pathway-specific parameter values yield diverse, ligand- and dose-specific deployment of NFκB signaling codewords.

### Oscillatory NFκB dynamics are a hallmark of paracrine TNF signaling

Overall, model simulations qualitatively matched measured trajectories at the respective doses. However, we identified a notable discrepancy in the responses of the MyD88-dependent pathway downstream of TLR9 at low doses (33 nM CpG, Fig. 5G). Simulations in this condition did not show substantial NFκB activation, but the measured trajectories showed oscillatory responses.

To address this discrepancy, we noticed that within the population of diverse responses to CpG, oscillatory trajectories were generally slightly delayed compared to transient and non-oscillatory trajectories (Fig. 6A). We therefore wondered whether cytokine feedback, especially by TNF ^41^, not represented in the simple mathematical models, might be responsible for this discrepancy between model simulations and experimental observations. Indeed, we found that a small but statistically significant amount of TNF was detectable in the cell culture medium at the early 5-minute timepoint of CpG stimulation (Fig. 6B). Furthermore, flow cytometry for the TNF receptor revealed a rapid internalization of TNFR1 not only in response to TNF but also CpG, which was also TNF-dependent (Fig. 6C). To test whether paracrine TNF signaling was in fact responsible for oscillatory NFκB responses, we measured single-cell dynamic responses to CpG in the presence or absence of saturating levels of recombinant soluble TNFRII (Fig. 6D). We noted a substantial decrease in oscillatory trajectories and the fraction of non-responding cells increased in the TNF blocking condition (Fig. 6D, 6E). Our data suggests that TNF produces oscillatory NFκB activity within cell populations exposed to low levels of CpG. We imagine that cells, which are unresponsive to CpG due to for example low TLR9 levels, may still respond to TNF produced by cells in the population that are responsive to CpG, possibly because of higher levels of TLR9 (Fig. 6F). Thus, in the context of MyD88-mediated PAMPs, oscillatory NFκB may be an indicator of paracrine signaling by host factor TNF.

**Figure 6.**
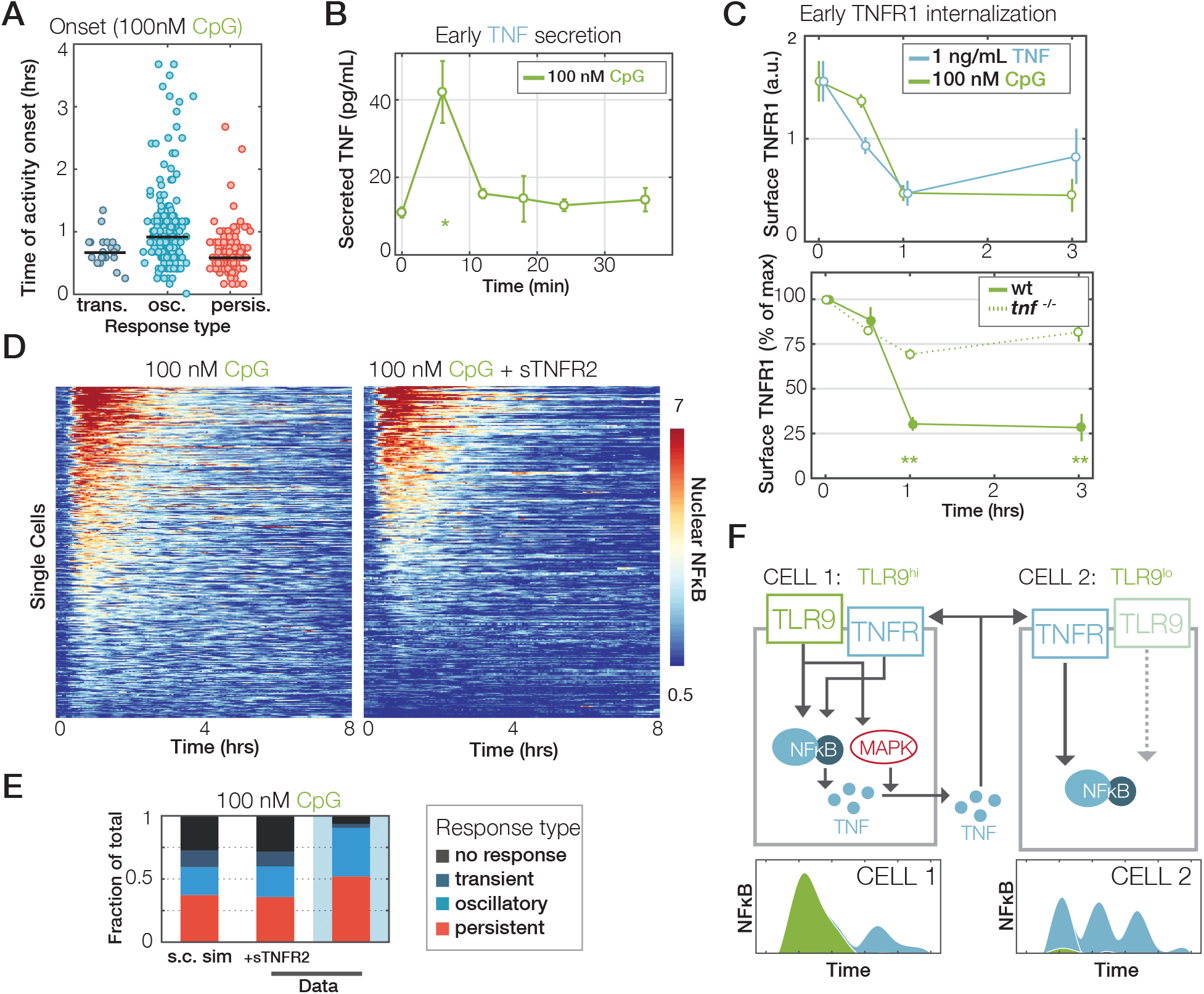
Oscillatory NFκB in response to PAMPs is a hallmark of feedforward TNF. (A) Activity onset times in single-cell NFκB responses to 100 nM CpG, grouped by dynamic subtypes of the response (persistent, oscillatory, or transient). (B) Early-phase TNF secretion dynamics from macrophages stimulated with 100 nM CpG, as measured by ELISA. (C) Top: median surface TNFR1 expression over time in BMDMs exposed to 1 ng/mL TNF or 100 nM CpG, monitored by flow cytometry. Bottom: median surface TNFR1 expression over time in wt or *Tnf*^−/−^ BMDMs in response to 100 nM CpG (scaled to receptor levels before treatment). Error bars show standard deviations across 3 independently-performed experiments, and double asterisks indicate a p-value <0.001 using a Student’s t-test comparing wild-type and knockout levels at a particular timepoint. (D) Single-cell heatmaps of NFκB activation mVenus-RelA BMDMs in response to 100 nM CpG, with or without feedforward TNF signaling blocked using saturating amounts (5 μg/mL) of soluble TNFRII co-injected with treatment. (E) Proportions of NFκB dynamic subtypes (off, transient, oscillatory, or persistent) as quantified from the data in (D). (F) Schematic depicting two cells; one cell (left) responds to CpG by activating NFkB and producing TNF that may act upon it in an autocrine manner. Another cell (right) does not respond to CpG (possibly because of low TLR9 expression), but responds to paracrine TNF and hence producing oscillatory NFκB activity.

## DISCUSSION

In this work, we report the identification of six dynamical features that characterize complex, stimulus-specific timecourse trajectories of NFκB activities in single primary macrophage cells. Using information-theoretic and machine learning approaches, we show that these function as signaling codewords to convey information about the extra-cellular environment to nuclear target genes. Strikingly, in an inflammatory disease mouse model, diminished ligand-specific deployment of two codewords – “oscillation” and “duration” – results in greater confusion of ligand sensing that may contribute to the etiology of autoimmune diseases. Our investigation of the molecular mechanisms underlying the stimulus-specific generation of NFκB signaling codewords revealed simple circuit motifs responsible for each; plus the recognition that NFκB oscillations observed in macrophages are in fact often a hallmark of paracrine TNF.

These finding were made possible by our development of new experimental and computational tools that provided an unprecedented quantity and quality of experimental data in primary cells responding to diverse immune threats. As cell lines show reduced responsiveness^42^, and ectopic expression of reporters can lead to artefactual oscillatory dynamics ^43^, we generated a new NFκB RelA-Venus knock-in mouse strain allowing us to image primary macrophages, the cell type that functions as the sentinels of the immune system. We were able to study not only the dose-response relationships but also characterize NFκB responses to pathogen-derived and host-derived ligands to which these primary macrophages respond vigorously. A robust automated image analysis pipeline and newly developed information-theoretic analysis and machine learning classification workflows enabled a rigorous, quantitative analysis of over 4.9 million single-cell data points derived from 44 distinct time-course conditions.

To identify signaling codewords, informative dynamical features, we employed an information theoretic framework. Previous applications of an information theoretic framework related the timeseries of nuclear NFκB abundance at either one or several timepoints to different doses of stimulus ^24,28^. While it was shown that timecourse measurements can provide more information about ligand and dose than a single timepoint, ^44^ it remained unclear which dynamical features are important in conveying this information. Prior studies sought to characterize temporal NFκB trajectories in terms of ad-hoc-defined dynamical features such “duration” ^45,46^ or “inter-peak time/frequency” ^26^. However, these features were not tested for information content, though some appear to correlate with gene expression responses ^47,48^. Our datasets and analytical workflow allowed for an unbiased evaluation of hundreds of potential features and yielded six specific dynamical features that essentially define stimulus-specific NFκB dynamics for the five ligands at multiple doses tested here. As the identified dynamic signaling features optimally provide the nucleus with information about the extra-cellular environment, they are codewords of a signaling code. We showed that signaling codewords identified by the information-theoretic approach are sufficient for a machine to learn to correctly classify NFκB trajectories in terms of stimulus and dose. Interestingly, “inter-peak time” or “period” were not represented, but instead, the presence or absence of oscillatory content emerged as an important “codeword” – it is key to distinguishing PAMP-responsive and cytokine TNF-responsive NFκB dynamics. It will be of interest to determine whether with additional datasets from macrophages or other cell types, additional codewords of the NFκB signaling code may be identified.

We have begun to characterize the key mechanisms that encode the six signaling codewords of the NFκB signaling code. Building upon prior mathematical models that have investigated NFκB dynamics in response to a single ligand in immortalized cell lines ^49^, a new model presented here recapitulates for the first time both oscillatory and non-oscillatory trajectories in primary macrophages in response to five ligands at several different doses and provides insights into the molecular mechanisms. As the IκB-NFκB signaling module is common to all stimulus-response pathways, and the IκBα negative feedback loop indeed supports both oscillatory and non-oscillatory activities (Fig. 5C), stimulus-specific deployment of the six signaling codewords depends on the biochemical characteristics of components in the receptor-associated signaling modules. Key characteristics are (i) the ligand half-life, as short half-lives (e.g. TNF) render the duration of the response dependent on stimulus-concentration^37,43^, (ii) the receptor translocation and replenishment rates that may either allow for post-stimulation shutdown or second phase signaling ^50^, (iii) the dose-response of the adaptor (TRAFs, MyD88, TRIF), as for example MyD88 tends to digitize responses, but TRIF does not^42^, and (iv) the de-activation kinetics of adaptors and ubiquitin chain networks that are likely key determinants of the termination of signaling but require further biochemical characterization.

It is well established that the temporal trajectories of NFkB activity are correlated with gene expression^45–48^. Our results suggest that the codewords identified here may specify the stimulus-specific expression of those genes that are able to differentiate between their relative presence or absence. Some prior work has described molecular mechanisms that particular target genes employ to “decode” specific NFκB signaling codewords. “Peak amplitude/fold change” for example was described to be sensed effectively by an incoherent feedforward loop involving the NFκB-responsive generation of p50 homodimers^51^. Stimulus-specific “Duration” was found to be differentiated by two mechanisms: whereas stimulus-specific expression of core regulators of the inflammatory response was mediated by an mRNA half-life of a few hours, pro-inflammatory initiators tend to employ a chromatin-based mechanism that involves the movement of a nucleosome ^52^. Interestingly, the “oscillatory/non-oscillatory” codeword does not necessarily control the stimulus-specific expression of primary response target genes ^43^, but appears to determine the capacity of NFκB to generate *de novo* enhancers in macrophages and thus affects the potential for gene expression induced by subsequent stimuli ^53^.

A hallmark of all single-cell datasets is the heterogeneity within an isogenic, identically stimulated population. Hence it is not surprising that the stimulus-specificity of the dynamical features identified here is by no means perfect, and that a machine learning classifier applied to all features or the six most informative “codewords” reveals some confusion, particularly among the NFκB responses to three bacterial PAMPs. Confusion here means for example that some (but not all) cells stimulated with CpG produces NFκB responses that are indistinguishable from some (but not all) cells stimulated with Pam3CSK4. We suggest that the capacity (or lack thereof, i.e. confusion) for mounting specific responses is a fundamental functional characteristic of macrophages as immune sentinel cells. Furthermore, given macrophage’s functional plasticity, we expect that this characteristic of capacity for stimulus-discrimination be similarly tuned—determined by the context of microenvironmental cytokines and exposure histories. In this study, macrophages derived from a mouse model of the systemic inflammatory disease, Sjögren’s syndrome, showed dramatically increased levels of ligand confusion. This particular model involves genetic variants in the IκBα promoter but the impact on NFκB signaling dynamics at the single cell level was unknown. While cells are capable of responding to diverse immune threats, the reduction in specificity adds to our understanding of this systemic autoimmune disease and may contribute to its etiology. Future studies will address whether these concepts apply to other autoimmune or inflammatory diseases.

Within the context of the innate immune signaling network, NFκB is only one of four prominent PAMP-responsive pathways, and therefore functions in conjunction with the JNK-AP1, MAPKp38/ERK-TTP/CREB, TBK-IRF3-ISGF3 axes to control gene expression responses ^54^. While bacterial ligands LPS, CpG, Pam3CSK4 are poorly distinguished at the level of NFκB dynamics (Fig. 4A), only LPS activates TBK-IRF3-ISGF3 and thus produces highly distinct gene expression program. Furthermore, MAPKp38 activation is specific to the high dose of LPS ^55^ allowing for much better dose distinction than that based on NFκB dynamics alone. Elucidating how coordinated combinatorial and temporal codes specify macrophage responses to diverse stimuli will be a major undertaking of future research.

## METHODS

### Experimental Methods

#### Mouse Models

The mVenus-RelA endogenously-tagged mouse line was generated by Ingenious Targeting Laboratory. A donor sequence encoding the monomeric variant of the Venus fluorescent protein ^56^ joined by a short flexible linker sequence directly upstream of the start codon of the murine *Rela* locus was used to generate, via homologous recombination, a tagged embryonic stem cell line, that was implanted to yield heterozygous mice. These mice were then bred with a mouse line constitutively expressing the *Flp* recombinase to remove the *Neo* resistance marker included in the homologous donor sequence. We then back-crossed the resultant mice with wild-type C57BL/6J mice to remove the *Flp* background and generate homozygously tagged mice (RelA^V/V^). mVenus-RelA mice were crossed into a IκBα^−/−^TNF^−/+^cRel^+/−^ line (TNF and cRel heterozygosity are required to rescue embryonic lethality of the IκBα^−/−^ genotype) ^57^, as well as into an IκBβ^−/−^ IκBε^−/−^ knockout line ^45^. For the Sjӧgren’s syndrome mouse model, we crossed mVenus-RelA mice into a strain that harbors mutated κB sites in the IκBα promoter ^6^.

#### Macrophage Cell Culture

Bone marrow-derived macrophages (BMDMs) were prepared by culturing bone marrow monocytes from femurs of 8-12 week old mice in CMG 14-12-conditioned medium using standard methods ^42,58^. BMDMs were re-plated in experimental dishes on day 4, then were stimulated on day 7. BMDMs were stimulated with indicated concentrations of lipopolysaccharide (LPS, Sigma Aldrich), murine TNF (R&D), a TLR1/2 agonist, the synthetic triacylated lipoprotein Pam3CSK4 (PAM), a TLR3 agonist, low molecular weight polyinosine-polycytidylic acid (Poly(I:C) (PIC)), a TLR9 agonist, the synthetic CpG ODN 1668 (CpG).

#### Biochemical assays

For immunoblots of whole cell lysates, bone-marrow derived macrophages were replated on day 4 at 20,000/cm^2^ in 6-cm dishes. After stimulation on day 7, sample buffer was added to 6-well plates directly after washing cells with PBS. Immunoblots followed standard procedure with anti-RelA (sc-372, Santa Cruz Biotechnology), anti-pIKK (CST2697), and IKK2 (CST2678). Western blot band intensities were quantified using ImageJ. Nuclear extract preparation and electrophoretic mobility shift assays followed published procedures^59^.

#### Live-cell imaging

Bone-marrow macrophages were replated on day 4 at 20,000 or 15,000/cm^2^ in an 8-well ibidi SlideTek chamber, for imaging at an appropriate density (approx. 60,000/cm^2^) on day 6 or day 7. 2 hours prior to stimulation, cells were incubated for 5 minutes at room temperature in a solution of 2.5 ng/mL Hoechst 33342 in PBS, then BMDM culture media was replaced. This staining condition was optimized to ensure no loss of cell viability and no aberrant morphological changes over a 24 hr period of imaging in the conditions described below. Cells were imaged at 5-minute intervals on a Zeiss Axio Observer platform with live-cell incubation, using epifluorescent excitation from a Sutter Lambda XL light source. Images were recorded on a Hamamatsu Orca Flash 2.0 CCD camera. After the start of imaging, additional culture media containing stimulus (TNF, LPS, poly(I:C), CpG, or Pam3CSK4) was injected into the chamber *in situ*. We have documented the reliability of the imaging workflow by establishing that distinct biological replicates give reproducible data (Supplementary Fig. 1D) and that distinct imaging frames of the same well provide reproducible data (Supplementary Fig. 1E). All data are available at https://data.mendeley.com/datasets/6wksmvh5p4/draft?a=832656ba-2bde-40a4-8bbc-4cecb1d9543d.

#### Measurement of TNF secretion (ELISA)

Bone-marrow macrophages were replated on day 4 at 25,000/cm^2^ in a 96-well format. On day 6, media was refreshed with 80 μL media containing indicated treatment (TNF, LPS, or CpG). Supernatants were collected from wells, in triplicate, at indicated timepoints, using procedures from the murine TNF alpha ELISA Ready-SET-Go! kit (eBioscience #88-7324-88). To optimize assay sensitivity, measurement was performed in a half-area 96-well plate (Corning #3690), and sample incubation was performed overnight at 4°C. Fluorescence measurements were performed using a standard spectrophotometer.

#### Measurement of surface receptor expression (FACS)

Bone-marrow-derived macrophages were replated on day 4 at 20,000/cm^2^ in 6-cm dishes. On day 6, media was refreshed with 3 ml media containing indicated treatment (TNF, LPS, or CpG). At indicated time point, media was rinsed out with cold PBS. Cells were incubated with fluorophore-conjugated antibodies for TNFR, CD11b, and F4/80 (BioLegend #113005, eBioscience #11-0112-82, eBioscience #12-4801-82) and analyzed, in triplicate. Antibody concentration and staining conditions were performed according to manufacturer recommendations. Stained cells were measured using an Accuri C6 Flow Cytometer (BD Biosystems). Fluorescence compensation and live/dead cell filtering was performed in FlowJo v10.

### Image analysis

Microscopy time-lapse images were exported for single-cell tracking and measurement in MATLAB R2016a. The tracking routines followed those used in earlier work ^44^. Briefly, cells were identified using DIC images, then segmented, guided by markers from the Hoechst image. Segmented cells were linked into trajectories across successive images, then nuclear and cytoplasmic boundaries were saved and used to define measurement regions in other fluorescent channels, including mVenus-NFκB. Nuclear NFκB levels were quantified on a per-cell basis, normalized to image background levels, then were baseline-subtracted. Mitotic cells, as well as cells that drifted out of the field of view, were excluded from analysis. The toolboxes used for this analysis are available at https://github.com/brookstaylorjr/MACKtrack.

### Channel capacity calculation and codeword identification

#### Feature selection

As there are ~ 9.3 × 10^16^ seven-dimensional combinations of 918 features (Supplementary Table 3) and each channel capacity calculation takes ~90 seconds per combination, evaluating channel capacity of all combinations of features would take ~2.3 × 10^15^ hours (~2.7 × 10^11^ years) to compute and is therefore is computationally infeasible. To narrow the search space, we utilized a feature selection approach. Since the channel capacities of individual features combine nonlinearly, there is no guarantee a high-ranking feature in low dimensional space will also be a subset of a high-ranking feature vector in high-dimensional space. Consequently, we utilized a forward feature selection approach that balances channel capacity rankings in lower dimensional space and diversity of candidates. Channel capacity calculations are performed on single dimensional features, ranked, and a subset of features above a threshold are selected to maximize diversity. As such 1D candidates are combined to form a set of 2D feature vectors. Channel capacity calculations are calculated on the 2D feature vectors, ranked and selected as in the 1D case. This iterative ranking and selection processes are repeated until additional dimensions offer no gain in channel capacity (Supplementary Table 4). Details are found in Supplementary Note 1.

### Machine Learning Classifier

#### Construction of classification models

We trained an ensemble of 100 decision trees using the fitcensemble function from the Statistics and Machine Learning Toolbox from MathWorks. Decision tree models are simple, highly interpretable, and can be displayed graphically ^60^. Consequently, the decision process of the classifier can be easily interrogated. However, decision tree models have two key disadvantages: (1) mediocre prediction performance ^61^ and (2) high variance due to overfitting ^60^. Both disadvantages can be mitigated by aggregating an ensemble of decision trees. Empirical comparison of classification models show that ensembles of decision trees outperform other classification algorithms across of a variety of problem sets ^61^.

We used a bootstrap aggregation (bag) method for constructing the ensembles. Each tree in the ensemble is trained on a bootstrapped replica of the data—each replica is a random selection of the data with replacement. The predictions from the ensemble model are determined by a majority vote from each individual tree prediction. We trained the ensemble to learn the stimulus labels (TNF, Pam3CSK4, CpG, LPS, and Poly(I:C)) from either the entire set of predictors (all 918 metrics, Supplementary Table 3) or a subset of predictors termed “signaling codewords” (Supplementary Table 3).

#### Decision tree parameters

To construct each decision tree, the software considers all possible ways to split the data into two nodes based on the values of every predictor. Then, it chooses the best splitting decision based on constraints imposed by training parameters, such as the minimum number of observations that must be present in a child node (MinLeafSize) and a predictor selection criterion. The software recursively splits each child node until a stopping criterion is reached. The stopping criteria include (1) obtaining a pure node that contains only observations from a single class, (2) reaching the minimum number of observations for a parent node (MinParentSize), (3) reaching a split that would produce a child node with fewer observations than MinLeafSize, and (4) reaching the maximum number of splits (MaxNumSplits). We used default values for MinLeafSize, MinParentSize, and MaxNumSplits: 1, 10, sample size – 1, respectively ^62^. Loadings flor classification models are listed in Supplementary Table S6.

Since the standard prediction selection process at each node may be biased, we used a predictor selection technique, interaction-curvature test, which minimizes predictor selection bias, enhances interpretation of the model, and facilitates inference of predictor importance. The interaction-curvature technique selects a predictor to split at each node based on the *p*-values of curvature and interaction tests. Whereas the curvature test examines the null hypothesis that the predictor and response variables are unassociated, the interaction test examines the null hypothesis that a pair of predictor variables and the response variable are unassociated. A node with no tests that yield *p*-values ≤ 0.05 is not split. At each node, the predictor or pair of predictors that yield the minimum significant *p*-value (0.05) is chosen for splitting. To split the node, the software chooses the splitting rule that maximizes the impurity gain—difference in the impurity of the node (calculated using Gini’s diversity index) and the impurity of its children nodes ^62^.

#### Evaluation

We evaluated the performance of the classifiers using 5-fold cross validation, an independent testing data set, or out of bag cross validation. We used the following performance metrics: true positive rate (recall), positive predictive value (precision), area under the Receiver Operating Characteristic (ROC) curve, F1 score, Matthews correlation coefficient, markedness, informedness and mean classification margin ^63–65^.

### Mathematical Modeling

#### Model structure

Several related models of NFκB activation in response to TNF have been established and iteratively parameterized ^45,66,67^, and used as a basis for modeling the NFκB response to LPS and other stimuli in immortalized cell lines with exogenously introduced (and overexpressed) fluorescent RelA ^42,68^. The model presented here to account for NFκB dynamics in primary macrophages is closely based on these previous studies, inheriting identical model topologies where possible and minimizing any changes to parameter values.

#### Key experimental data constraints

As a first step towards parameterizing our model, we quantified characteristics of oscillatory endogenous BMDM signaling. We observed only slight differences in peak periodicity and amplitude between conditions (roughly a 10-minute difference in median period for the lowest dose of TNF which induced robust oscillations, 0.33 ng/mL, and the highest dose tested). We did, however, observe pronounced differences in duration as the dose of TNF is increased (Supplementary Figure 1F). Median period was determined to generally fall within 90-95 min, in the same range of oscillations measured in other cell types ^66,67^.

These oscillations appeared to be remarkably stable across an extremely broad range of induction levels. Indeed, the variation observed across single cells in a particular condition (or even within the same cell) is much smaller than any differences in oscillations observed between conditions. Even when other stimuli are considered, the “signature” first harmonic of the oscillatory subpopulation remains consistent. This consistency across a wide range of input conditions agrees, notably, with predictions made using simplified discrete delay model of the NFκB network ^69^. These delays could plausibly arise from IκB mRNA (measured to be some 10-12 minutes) ^70^ and protein processing.

Biochemical assays indicate that the major difference between TNF and LPS-induced IKK activation is not in the maximum amplitude, but the duration of IKK induction ^57,71^. TNF strongly but transiently activates IKK. Peak IKK activity is limited in duration by rapid internalization and degradation of the ligand-bound receptor ^72–74^. LPS-bound TLR4 is also rapidly internalized, but continues to strongly activate IKK from the endosome ^75^. This difference is reflected in single-cell NFκB activation: while the speed of NFκB activation (roughly proportional to the peak of IKK activity) is similar between TNF and LPS, sustained high levels of IKK activity in response to LPS leads to higher peak activity (Supplementary Figure 5B, C).

#### Model Fitting

We first sought to fit models for TNFR signaling and TLR4 signaling. We employed a screen where repeated, random initialization of parameters was followed by their optimization, fitting model predictions to IKK activation dynamics established by kinase assay measurements ^42,76^. The resultant model required rapid IKK de- and re-activation ^77^, which allowed IKK responses to be both adaptive (in the case of TNF) and long duration (as in TLR4 responses).

For the IKK-IκB-NFκB core module, model topology and parameters were confined to be near previously established values (Supplementary Table 7). We performed a multidimensional sweep of transport rates and found a narrow range of parameters that could account for the observed frequency invariance, with high IKK activity diminishing oscillatory behavior (Supplementary Figure 5D). A subsequent fitting process allowed us to optimize other parameters, including the induced synthesis rate constant of IκBα and the activation rate constant of IKK. We repeated this two-stage sweep/fitting process until parameter values converged.

For signaling via TLR1/2, TLR3, and TLR9, we used prior estimates of each receptor’s abundance in monocytes/macrophages ^40^ to estimate synthesis and degradation rates. In many cases, receptor-ligand affinities were also known ^78–80^ and were therefore used to estimate association and dissociation of the receptor. The kinetics of each receptor’s association with a downstream adaptor (TRIF or MyD88) were taken from estimates from our TLR4 model. NFκB responses to TLR9 were observed to be more transient than to either TLR4 or TLR1/2, in agreement with previous data ^59^ and the observed self-inactivation of TLR9 ^81^.

The software to run the model is available at https://github.com/Adewunmi91/nfkb_model.

## DATA AND SOFTWARE AVAILABILITY

Data is available in https://data.mendeley.com/datasets/6wksmvh5p4/draft?a=832656ba-2bde-40a4-8bbc-4cecb1d9543d. Software associated with the image processing and computational modeling are available at https://github.com/brookstaylorjr/MACKtrack, https://github.com/Adewunmi91/nfkb_model.

## ACKNOWLEDGEMENTS

We thank laboratory members, especially Quen Cheng and Ying Tang, as well as Roy Wollman (UCLA) and Eric Deeds (UCLA), for critical discussion and reading of the manuscript. The work was supported by NIH MSTP and Vascular Biology training grants, and an NIH NRSA Predoctoral Fellowship to A.A. (T32GM008042, T32HL69766, F31AI138450), a postdoctoral fellowship from the Deutsche Forschungsgemeinschaft (DFG, German Research Foundation, 419234150) to S.L., and NIH grants to A.H. (R01GM117134, R01AI127864).

## AUTHOR CONTRIBUTIONS

B.T. designed the knockin mouse, performed the cell imaging, developed the image analysis workflow, the information theoretic workflow, and the mathematical model. A.A. performed the NFκB EMSA, the machine learning analysis and the experimental and analytical work with the Sjögren’s mouse model. Y.L performed the flow cytometry experiments. S.L. performed IKK immunoblot analysis. B.T., A.A., and A.H. wrote the paper, all authors proofed the paper.

