## Supplementary Figures 1-6, Tables 2-4, Notes for "Identification and physiological significance of temporal NFκB signaling codewords deployed by macrophages to classify immune threats"

**This PDF file includes:**

**Supplementary Figures 1 to 6, related to Figures 1 to 6.**

**Supplementary Tables 2 to 4.**

**Supplementary Note 1.**

**Other Supplementary Information for this manuscript, not included in this PDF:**

**Supplementary Table 1**
Compilation of the literature on NFκB dynamic control.

**Supplementary Table 5.**
Specification of the violin plots of Figure 3A.

**Supplementary Table 6.**
Loadings of machine learning models of Figure 3B/D/4A (ligand prediction) and 3C/E (dose prediction), and Figure 4E. All values were L1 normalized so that the sum of all absolute quantities equals 1.

**Supplementary Table 7.**
Specification of the Mathematical Model: reactions and parameter values. Color-code indicates the regulatory module. Reaction 1-26, 67 (black): common core module; reaction 27-52 (red): TLR4 module; reaction 53-66 (blue): TNFR module; reaction 68-76 (orange): TLR2 module; reaction 77-84 (purple): TLR3 module; reaction 85-94 (green): TLR9 module.

**Supplementary Movie 1.**
Distinct cellular responses to TNF and LPS. Live cell imaging and automated image analysis to quantify the NFκB trajectory.

**Data 1.**
Online Data folder including all live cell imaging movies and derived data files.
https://data.mendeley.com/datasets/6wksmvh5p4/draft?a=832656ba-2bde-40a4-8bbc-4cecb1d9543d

**Software 1.**
Image analysis software is available at https://github.com/brookstaylorjr/MACKtrack .

**Software 2.**
Mathematical Modeling software is available at https://github.com/Adewunmi91/nfkb_model .

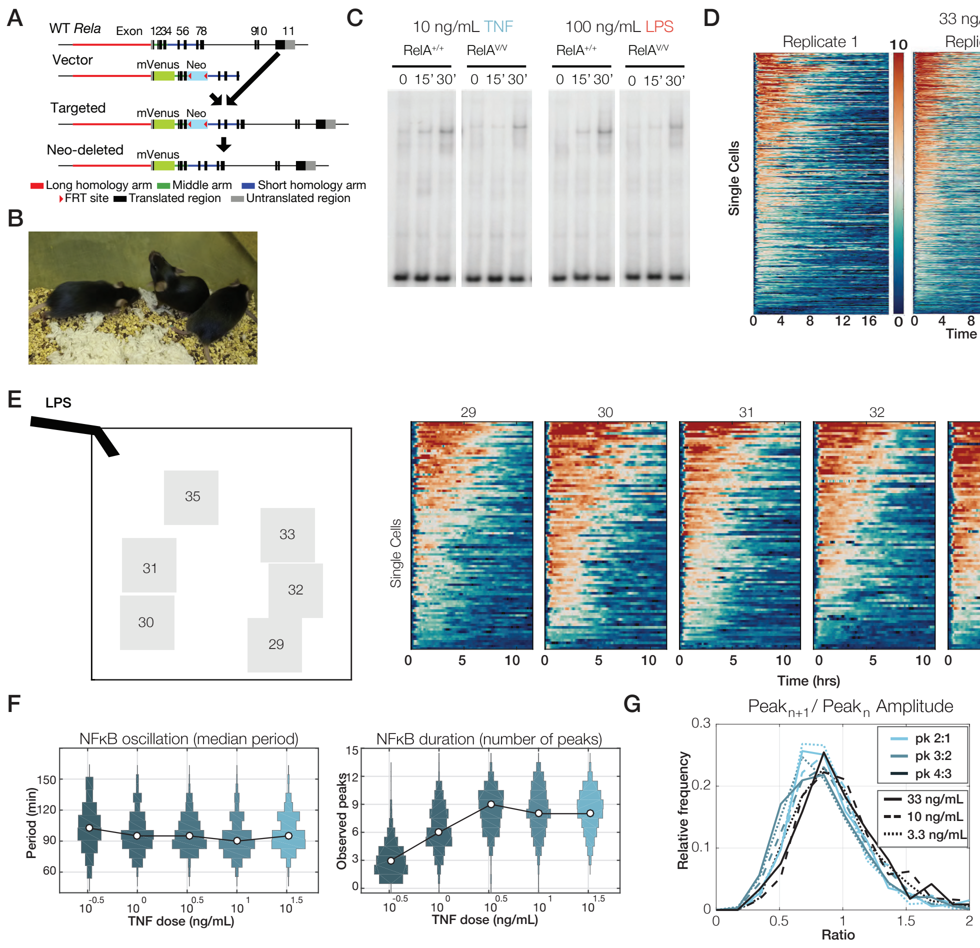

**Supplementary Figure 1. An experimental model and imaging workflow allows for reliable tracking NFκB RelA dynamics in primary macrophages at single cell level.**

(A) Schematic of the homologous recombination strategy for generating the mVenus-RelA allele in embryonic stem cells. These were injected into blastocysts for knockin mouse generation.

(B) Image of homozygous mVenus-RelA knockin mice shows that they are overtly healthy.

(C) mVenus-RelA macrophages show normal levels of NFκB DNA binding activity. NFκB EMSA of nuclear extracts made from mVenus-RelA and wild-type control BMDMs stimulated for 0, 15’, and 30’ with 10 ng/mL TNF and 100 ng/mL LPS.

(D) Experimental live cell imaging workflow and automated image analysis is robust as documented by biological replicates produced months apart from different mice.

(E) Microscopy workflow shows no location bias. Fields of view used in replicate 2 of 33 ng/mL LPS condition (left). Heatmaps of NFκB responses of cells in different fields of view (right).

(F) TNF dose does not regulate oscillation period but duration. Violin plots showing distributions of single-cell oscillation period (median peak-to-peak time) and duration (number of peaks measured in 18 hrs) across a range of TNF stimulus levels.

(G) TNF oscillations do not have a primary first peak, but rather steadily diminishing peaks. Histograms of oscillatory peak ratios (i.e. between amplitudes of subsequent peaks in the same cell) in response to multiple doses of TNF.

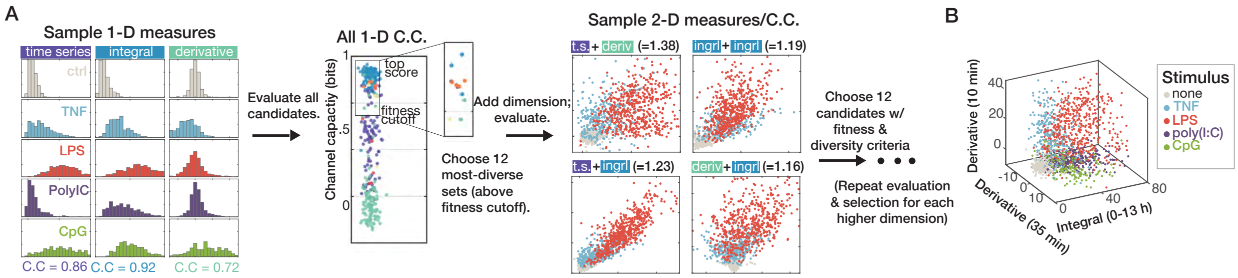

**Supplementary Figure 2. Design of an algorithm to identify information-maximizing combinations of dynamic features.**

(A) Procedure: single-dimension measurements (shown as histograms of cell population for each input condition) are evaluated across all input conditions. The output channel capacities are ranked: a subset of the candidates above a minimum "fitness" threshold are then selected to maximize diversity. These candidates are then re-evaluated in conjunction with a second dimension. This ranking/selection process is repeated until the final dimension is reached and a multidimensional vector is assembled.

(B) A sample representation of an optimal three-dimensional vector capturing single-cell measurements of NFκB responses. Using a three-dimensional vector, NFκB responses are quantified and each cell responding to the indicated ligand is depicted in a three-dimensional graph.

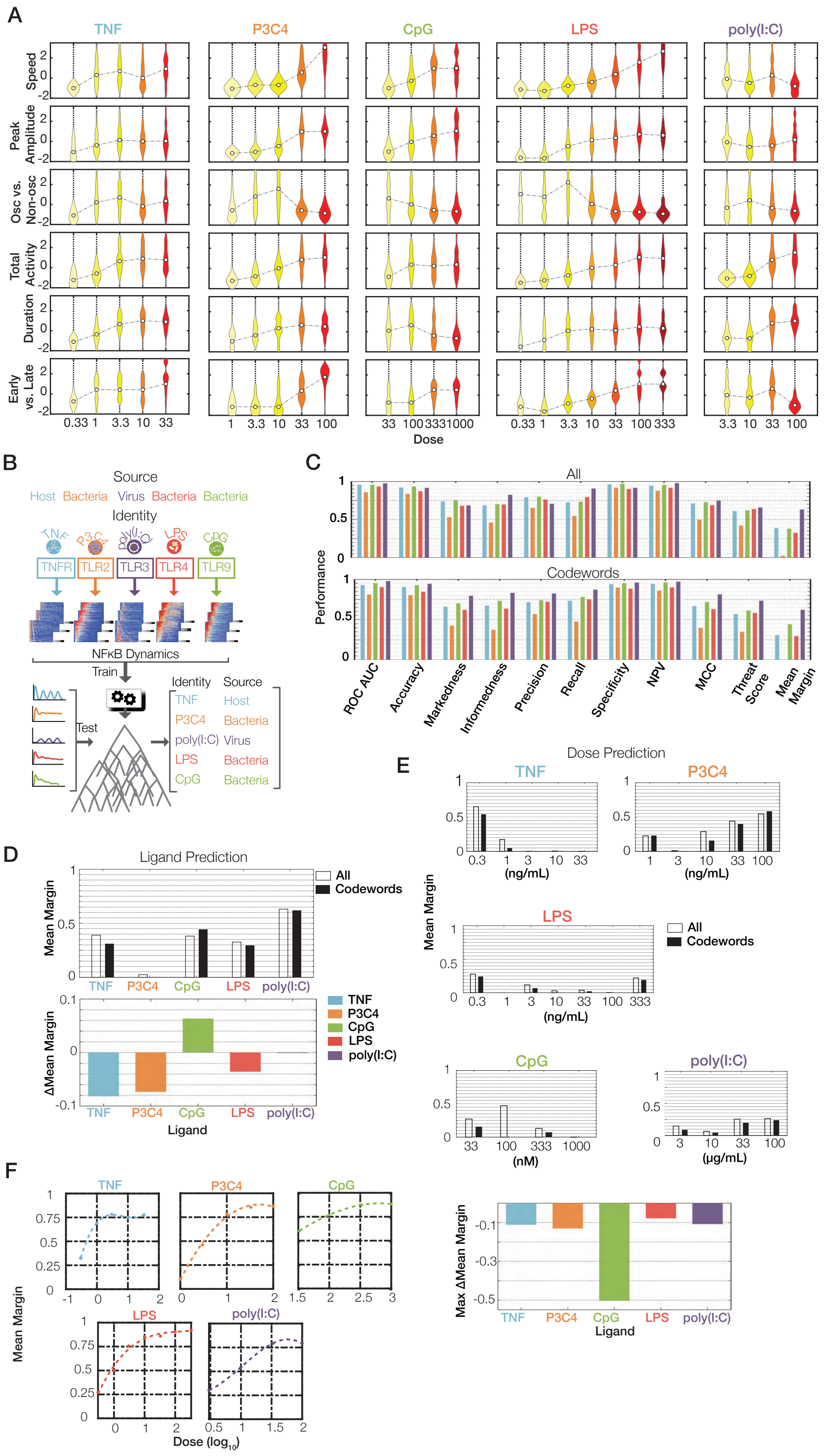

A
TNF
P3C4
CpG
LPS
poly(I:C)
Speed
Peak Amplitude
Osc vs. Non-osc
Total Activity
Duration
Early vs. Late
Dose
B
Source
Host Bacteria Virus Bacteria Bacteria
Identity
TNFR TLR2 TLR3 TLR4 TLR9
NFκB Dynamics
Train
Test
Identity Source
TNF Host
P3C4 Bacteria
poly(I:C) Virus
LPS Bacteria
CpG Bacteria
C
All
Codewords
Performance
ROC AUC
Accuracy
Markedness
Informedness
Precision
Recall
Specificity
NPV
MCC
Threat Score
Mean Margin
D
Ligand Prediction
Mean Margin
All
Codewords
ΔMean Margin
Ligand
E
Dose Prediction
(ng/mL)
(nM)
(μg/mL)
Max ΔMean Margin
F
Mean Margin
Dose (log10)

**Supplementary Figure 3. Evaluating a machine learning classifier: sufficiency of signaling codewords.**

(A) Dose-dependent deployment of signaling codewords. Violin plots of the relative presence of indicated codewords (z-score) in the trajectories of individual cells. Violin plots are specified in Supplementary Table 5.

(B) Schematic of the machine learning classification procedure: Predictors/features of NFκB signaling dynamics in response to TNF, Pam3CKS4, poly(I:C), LPS, and CpG were used to train an ensemble (using bootstrap aggregation) of 100 decision tree models to predict ligand identity and ligand source.

(C) Six codewords perform as well as all 918 dynamical features. A variety of performance metrics to ascertain the performance of ligand classifiers trained with all features (top) and or only codewords (bottom). Classifiers were evaluated using 5-fold cross validation or out-of-bag crossvalidation.

(D) Comparison of ligand classification margin (difference of the true ligand probability and maximum false ligand probability) of models trained using all predictors versus signaling codewords: (top) mean classification margins across all ligands; (bottom) difference of mean classification margin of codeword classifier and all predictors classifier.

(E) Comparison of dose classification margins of classifiers trained on all predictors versus codewords: mean classification margins across all doses for each ligand (top). Maximum of the differences in mean classification margins between codewords classifiers and all predictor classifiers (bottom).

(F) Dose dependence of ligand identification: mean margin of binary classifiers that distinguish no treatment controls from each ligand at the indicated dose.

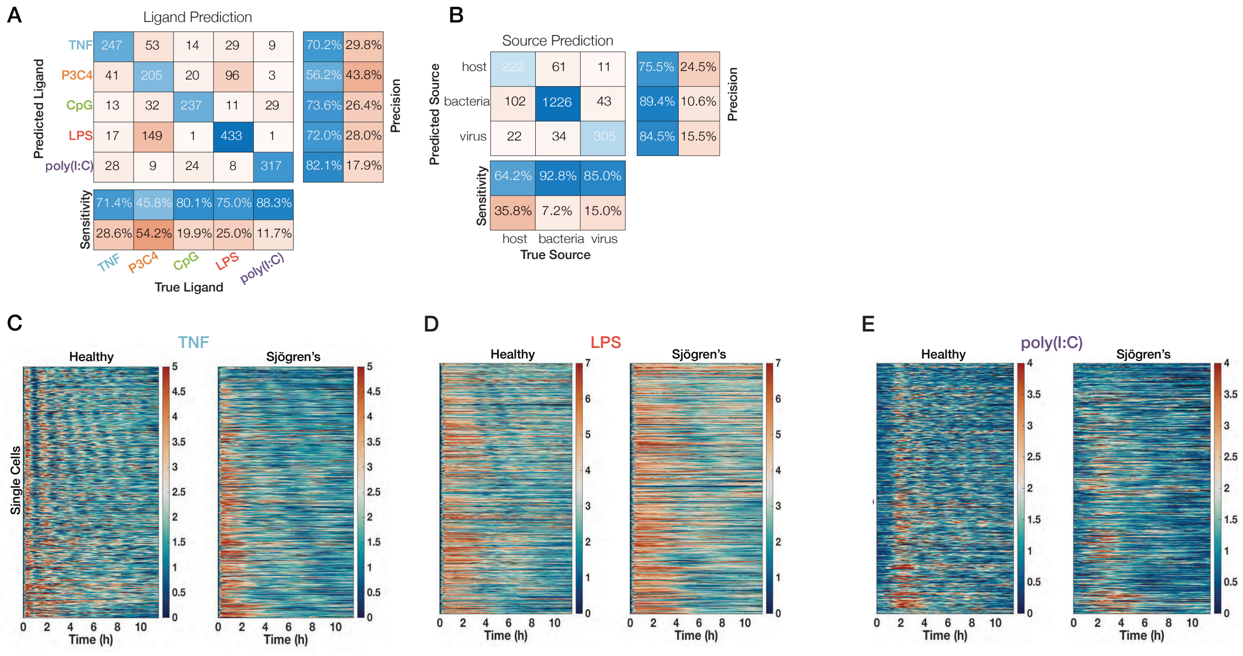

**Supplementary Figure 4. Analyzing stimulus confusion in macrophages from healthy and diseased mice.**

(A) Confusion matrix of ligand predictions: diagonal values show correct predictions and off-diagonals values show incorrect predictions; (right) percentage of correct predictions (precision; in blue) and incorrect predictions (false discovery rate; in orange); (bottom) percentage of ligands correctly identified (sensitivity/recall; in blue) and not identified (miss rate/false negative rate; in orange).

(B) Confusion matrix of source predictions:(bottom) sensitivity, (right) precision.

(C-E) Heatmaps of nuclear NFκB trajectories in hundreds of macrophages from healthy mice compared to Sjögren's mice in response to 10 ng/mL TNF (C), 100 ng/mL LPS (D), and 50 μg/mL poly(I:C) (E).

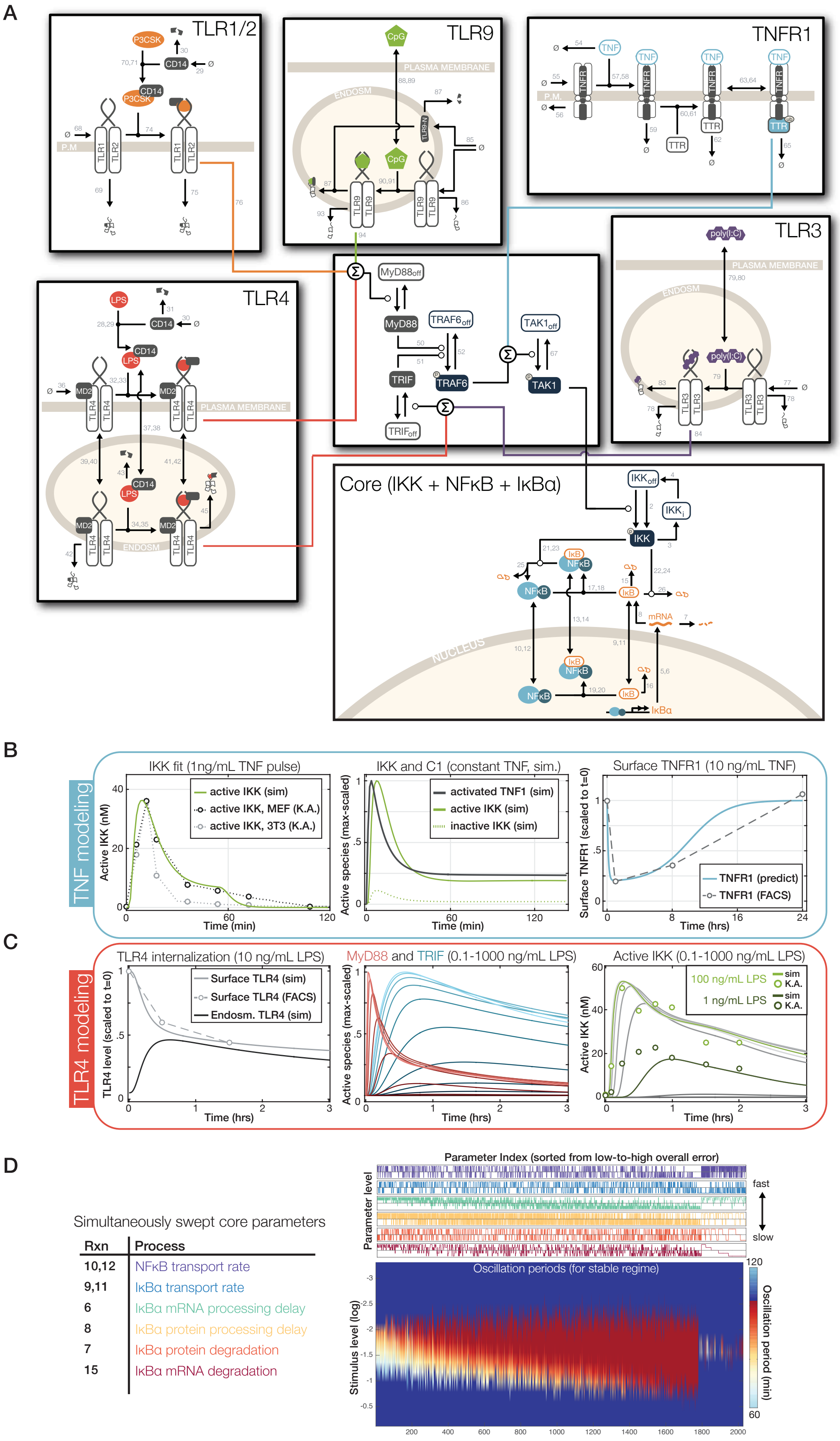
A
TLR1/2
P3CSK
CD14
TLR1
TLR2
P.M.
TLR9
CpG
PLASMA MEMBRANE
ENDOSM
TLR9
TNFR1
TNF
TNFR
TTR
P.M.
TLR4
LPS
CD14
MD2
TLR4
PLASMA MEMBRANE
ENDOSM
MyD88off
MyD88
TRAF6off
TRAF6
TAK1off
TAK1
TRIF
TRIFoff
TLR3
poly(I:C)
PLASMA MEMBRANE
ENDOSM
TLR3
Core (IKK + NFκB + IκBα)
IKKoff
IKKi
IKK
IκB
NFκB
mRNA
NUCLEUS
IκBα
B
TNF modeling
IKK fit (1ng/mL TNF pulse)
active IKK (sim)
active IKK, MEF (K.A.)
active IKK, 3T3 (K.A.)
Active IKK (nM)
Time (min)
IKK and C1 (constant TNF, sim.)
activated TNF1 (sim)
active IKK (sim)
inactive IKK (sim)
Active species (max-scaled)
Time (min)
Surface TNFR1 (10 ng/mL TNF)
TNFR1 (predict)
TNFR1 (FACS)
Surface TNFR1 (scaled to t=0)
Time (hrs)
C
TLR4 modeling
TLR4 internalization (10 ng/mL LPS)
Surface TLR4 (sim)
Surface TLR4 (FACS)
Endosm. TLR4 (sim)
TLR4 level (scaled to t=0)
Time (hrs)
MyD88 and TRIF (0.1-1000 ng/mL LPS)
Active species (max-scaled)
Time (hrs)
Active IKK (0.1-1000 ng/mL LPS)
100 ng/mL LPS
1 ng/mL LPS
sim
K.A.
Active IKK (nM)
Time (hrs)
D
Simultaneously swept core parameters
Rxn | Process
10,12 | NFκB transport rate
9,11 | IκBα transport rate
6 | IκBα mRNA processing delay
8 | IκBα protein processing delay
7 | IκBα protein degradation
15 | IκBα mRNA degradation
Parameter Index (sorted from low-to-high overall error)
Parameter level
fast
slow
Oscillation periods (for stable regime)
Stimulus level (log)
Oscillation period (min)

**Supplementary Figure 5. Constructing and fitting a mathematical model of NFκB activation dynamics in response to multiple stimuli.**

(A) Reaction schema for the multi-stimulus model of NFκB activation. Each box represents a regulatory module, with receptor-associated modules connecting into common core modules. All reactions are shown with numbers representing kinetic rate constants identified in Table S6.

(B) TNFR-associated module: Left: IKK activation in response to a 45 min pulse of TNF activation were fit using a screen where repeated, random initialization was followed by optimization: the best fit model of 1024 trials is shown. Middle: dynamics of TNFR1 and IKK activation. The transience of IKK activation is likely to be driven by rapid receptor internalization, not IKK inactivation, as has been previously hypothesized. Right: predicted levels of TNFR1 internalization in response to 10 ng/mL TNF, and measured surface TNFR1 levels as measured by FACS.

(C) TLR4-associated module: Left: dynamics of TLR4 internalization. TLR4 is also rapidly internalized in response to binding of LPS, but stays active in the early endosome. Middle: dynamics of MyD88 and TRIF activation in response to 0.1 (dark curves) to 1000 (bright curves) ng/mL LPS. Right: fitted levels of IKK activation (note that peak activation is only app. 25% greater than activity induced by TNF) in response to 0.1 (dark curves) to 1000 (bright curves) ng/mL LPS. Simulated and measured (by kinase assay) dynamics at 1 ng/mL and 100 ng/mL LPS are highlighted in green.

(D) Results of a simultaneous parameter sweep in the "core" NFκB model (IKK, NFκB, and IκBα). Swept parameters are indicated in table on left. 1728 out of 2000 parameter combinations showed activation in response to a range of IKK values. The full dose response was measured and ranked along the oscillatory characteristics to ensure that parameter sets are robust in this key characteristic.

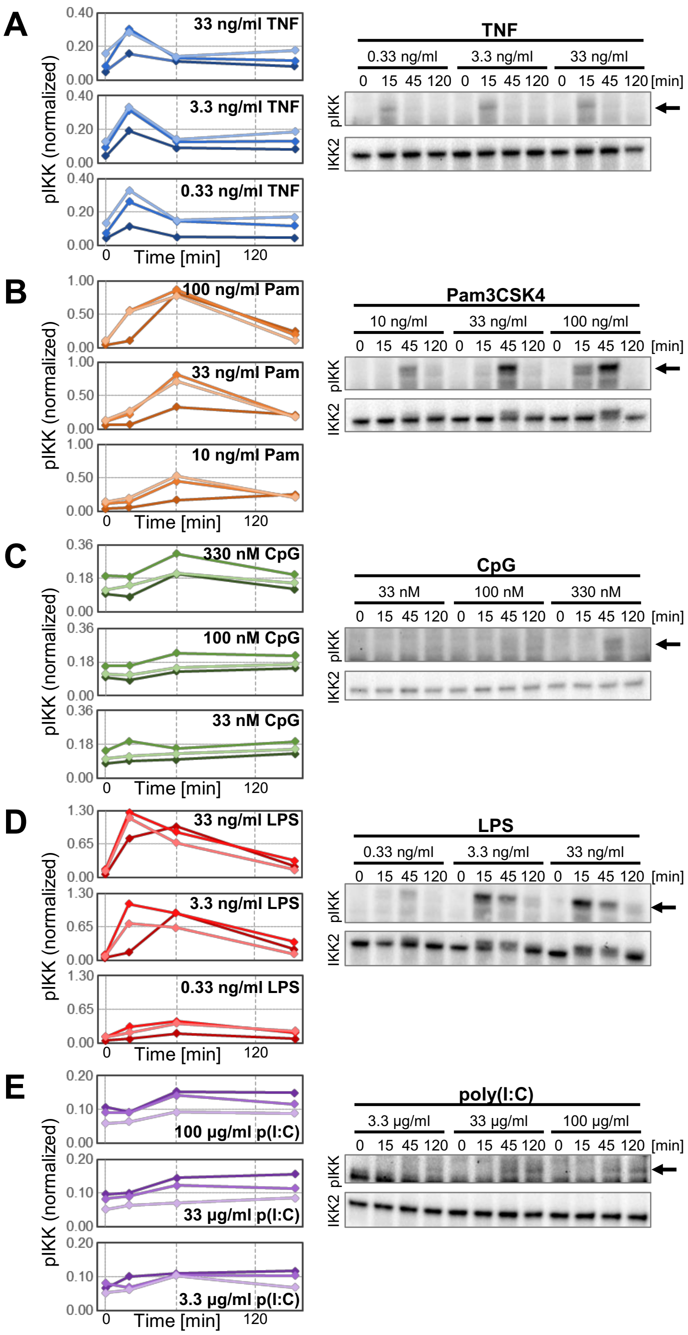

**Supplementary Figure 6. IKK activation dynamics in primary macrophages**

Levels of phosphorylated IKK (pIKK) in lysates of BMDMs stimulated with (A) TNF, (B) Pam3CSK4, (C) CpG, (D) LPS, and (E) poly(I:C) (HMW) at the indicated doses for 0, 15, 45, and 120 min were measured by immunoblotting after SDS-PAGE. For quantification, pIKK band intensities were normalized to total IKK2 levels and to a control stimulation (33 ng/ml LPS, 45 min, from replicate 1). (Left) Quantifications from three replicates are shown (dark shade: replicate 1, medium shade: replicate 2, light shade: replicate 3). (Right) Immunoblot from one representative experiment is shown. Arrow: pIKK band.

**Supplementary Table 2 – List of experiments, NFκB trajectories, data points**

| ID | Experiment Name | cell count | # of frames | # of datapoints |
|---|---|---|---|---|
| 274 | Ctrl | 229 | 259 | 51274 |
| 282 | 33 nM CpG | 347 | 259 | 85101 |
| 283 | 100 nM CpG | 392 | 259 | 98262 |
| 284 | 333 nM CpG | 423 | 259 | 107752 |
| 285 | 1 μM CpG | 296 | 259 | 74549 |
| 289 | 1 ng/mL LPS | 301 | 259 | 69536 |
| 290 | 0.33 ng/mL TNF | 488 | 259 | 119215 |
| 291 | 1 ng/mL TNF | 416 | 259 | 102095 |
| 292 | 3.3 ng/mL TNF | 433 | 259 | 106838 |
| 293 | 10 ng/mL TNF | 423 | 259 | 103614 |
| 294 | 33 ng/mL TNF | 346 | 259 | 85453 |
| 295 | 10 μg/mL Poly (I:C) | 370 | 259 | 87751 |
| 296 | 33 μg/mL Poly (I:C) | 423 | 259 | 103792 |
| 298 | 100 ng/mL Pam3CSK4 | 448 | 259 | 113463 |
| 300 | 0.33 ng/mL LPS | 458 | 259 | 106871 |
| 302 | 3.33 ng/mL LPS | 449 | 259 | 110748 |
| 303 | 10 ng/mL LPS | 617 | 259 | 155766 |
| 304 | 33 ng/mL LPS | 579 | 259 | 146270 |
| 305 | 3.3 μg/mL Poly (I:C) | 360 | 259 | 88191 |
| 306 | 100 μg/mL Poly (I:C) | 359 | 259 | 88831 |
| 309 | 33 ng/mL Pam3CSK4 | 467 | 259 | 116836 |
| 310 | 333 ng/mL LPS | 577 | 249 | 142330 |
| 311 | 100 ng/mL LPS | 546 | 249 | 134378 |
| 325 | 3.3 ng/mL Pam3CSK4 | 375 | 259 | 88425 |
| 326 | 1 ng/mL Pam3CSK4 | 287 | 259 | 66754 |
| 327 | 10 ng/mL Pam3CSK4 | 354 | 259 | 83585 |
| 335 | 33 ng/mL LPS | 532 | 259 | 135403 |
| 341 | 33 ng/mL LPS | 318 | 258 | 79633 |
| 352 | 33 ng/mL LPS | 590 | 259 | 150177 |
| **Subtotal Total** | | **12203** | | **3002893** |
| 367 | IκBα-/- : 3.3 ng/mL TNF | 570 | 224 | 114208 |
| 371 | IκBβ-/-ε-/-: 3.3 ng/mL TNF | 361 | 259 | 71956 |
| 374 | IκBβ-/-ε-/-: 10 ng/mL LPS | 455 | 259 | 103987 |
| **Subtotal Total** | | **1386** | | **290151** |
| 751 | IκBαM/M: 100 ng/mL LPS | 687 | 217 | 146420 |
| 752 | IκBαM/M: 50 μg/mL Poly I:C | 550 | 217 | 113094 |
| 753 | IκBαM/M: 10 ng/mL TNF | 632 | 217 | 130608 |
| 754 | 10 ng/mL TNF | 587 | 217 | 121493 |
| 755 | 50 μg/mL Poly I:C | 612 | 217 | 126201 |
| 756 | 100 ng/mL LPS | 731 | 217 | 154528 |

| | | | | |
|---|---|---|---|---|
| 759 | IκBαM/M: 100 ng/mL LPS | 733 | 217 | 156677 |
| 760 | IκBαM/M: 50 μg/mL Poly I:C | 500 | 217 | 103689 |
| 761 | IκBαM/M: 10 ng/mL TNF | 535 | 217 | 106993 |
| 762 | 10 ng/mL TNF | 462 | 217 | 91969 |
| 764 | 100 ng/mL LPS | 440 | 211 | 89399 |
| **Subtotal Total** | | **6469** | | **1341071** |
| **Total** | | **21444** | | **4924266** |

### Supplementary Table S3 – List of metrics used for information theoretic analysis

| Metric | Description | Parameter |
|---|---|---|
| Time Series | Nuclear NFκB level | time: 0-18 hrs, in 5 minute increments (217 values) |
| Integral | Accumulated NFκB activity | time: 0-18 hrs, in 5 minute increments (217 values) |
| Derivative | Central-difference estimate of instantaneous change in NFκB activity | time: 0-18 hrs, in 5 minute increments (217 values) |
| Maximum amplitude | Peak NFκB value | |
| Maximum integral | Peak level of accumulated NFκB | |
| Maximum derivative | Largest instananeous increase in NFκB | |
| Minimum derivative | Largest instantaneous decrease in NFκB | |
| Peak frequency | Frequency of 1st harmonic in NFκB activity | |
| Oscillatory component | Fraction of total signal energy that is high-frequency | Baseline "oscillatory" frequency spans 0.375 $hrs^{-1}$ to 0.625 $hrs^{-1}$ ) (8 values) |
| Peak 1 time | Timing of 1st recorded peak | |
| Peak 1 amplitude | Amplitude of 1st recorded peak | |
| Peak 2 time | Timing of 2nd recorded peak | |
| Peak 2 amplitude | Amplitude of 2nd recorded peak | |
| Consecutive duration | Maximum consecutive time NFκB activity was above baseline | Baseline activation value (spans 0-5.5 [roughly 0-75% of observed maximal activation level]) (25 values) |
| Duration | Total time NFκB activity was above baseline | Baseline activation value (spans 0-5.5 [roughly 0-75% of observed maximal activation level]) (25 values) |
| Integral (1-hour window) | Integrated activity within a defined window of time (e.g. 6-7 hrs) | Starting point (0-17 hrs in 0.5 hr increments) |
| Integral (3-hour window) | Integrated activity within a defined window of time (e.g. 6-9 hrs) | Starting point (0-15 hrs in 0.5 hr increments) |

**Supplementary Table 4A: Channel capacity measurements & metrics in optimized vectors across all ligands and doses**

| Dim | Metrics | Channel Capacity |
|---|---|---|
| 1 | integral_at_3.92h | 0.96 |
| 2 | amplitude_at_1.25h, derivative_at_0.08h | 1.38 |
| 3 | amplitude_at_1.25h, derivative_at_0.08h, duration_above_1.03 | 1.66 |
| 4 | amplitude_at_1.25h, derivative_at_0.08h, duration_above_1.03, amplitude_at_0.83h | 1.78 |
| 5 | derivative_at_0.5h, integral_at_13.25h, derivative_at_0.33h, derivative_at_0.75h, derivative_at_0.08h | 1.89 |
| 6 | derivative_at_0.5h, integral_at_13.25h, derivative_at_0.33h, derivative_at_0.75h, derivative_at_0.08h, duration_above_0 | 1.97 |
| 7 | derivative_at_0.5h, integral_at_13.25h, derivative_at_0.33h, derivative_at_0.75h, derivative_at_0.08h, duration_above_0, integral_at_7.75h | 2.02 |
| 8 | derivative_at_0.5h, integral_at_13.25h, derivative_at_0.33h, derivative_at_0.75h, derivative_at_0.08h, duration_above_0, integral_at_7.75h, minimum_derivative | 2.03 |
| 9 | derivative_at_0.5h, integral_at_13.25h, derivative_at_0.33h, derivative_at_0.75h, derivative_at_0.08h, duration_above_0, integral_at_7.75h, minimum_derivative, integral_at_18.58h | 2.05 |

**Supplementary Table 4B: Channel capacity measurements & metrics in optimized vectors across doses of TNF**

| Dim | Metrics | Channel Capacity |
|---|---|---|
| 1 | 1st_peak_amplitude | 0.68 |
| 2 | amplitude_at_0.25h, amplitude_at_0.75h | 0.79 |
| 3 | amplitude_at_0.25h, amplitude_at_0.75h, integral_at_17.92h | 0.90 |
| 4 | amplitude_at_0.25h, amplitude_at_0.75h, integral_at_17.92h, amplitude_at_7.17h | 0.96 |
| 5 | amplitude_at_0.25h, amplitude_at_0.75h, integral_at_17.92h, amplitude_at_7.17h, amplitude_at_7.83h | 1.00 |
| 6 | amplitude_at_0.25h, amplitude_at_0.75h, integral_at_17.92h, amplitude_at_7.17h, amplitude_at_7.83h, envelope_above_4.54 | 1.02 |
| 7 | integral_at_9.25h, integral_at_19.25h, envelope_above_0.62, derivative_at_0.25h, derivative_at_0.08h, duration_above_0.82, derivative_at_0.42h | 1.04 |

**Supplementary Table 4C: Channel capacity measurements & metrics in optimized vectors across doses of Pam3CSK4**

| Dim | Metrics | Channel Capacity |
|---|---|---|
| 1 | integral_at_1.08h | 0.85 |
| 2 | integral_at_4.75h, integral_at_1.25h | 1.26 |
| 3 | integral_at_4.75h, integral_at_1.25h, amplitude_at_0.33h | 1.41 |
| 4 | integral_at_4.75h, integral_at_1.25h, amplitude_at_0.33h, amplitude_at_1.75h | 1.46 |
| 5 | integral_at_4.75h, integral_at_1.25h, amplitude_at_0.33h, amplitude_at_1.75h, amplitude_at_0.75h | 1.48 |
| 6 | integral_at_4.75h, integral_at_1.25h, amplitude_at_0.33h, amplitude_at_1.75h, amplitude_at_0.75h, amplitude_at_2.5h | 1.51 |
| 7 | integral_at_2.58h, amplitude_at_0.25h, amplitude_at_0.33h, amplitude_at_2.42h, amplitude_at_0.83h, envelope_above_1.03, integral_at_8.58h | 1.52 |

**Supplementary Table 4D: Channel capacity measurements & metrics in optimized vectors across doses of PolyI:C**

| Dim | Metrics | Channel Capacity |
|---|---|---|
| 1 | integral_at_19.08h | 0.81 |
| 2 | integral_at_19.08h, amplitude_at_1.5h | 0.93 |
| 3 | integral_at_19.08h, amplitude_at_1.5h, amplitude_at_0.58h | 0.99 |
| 4 | integral_at_19.08h, amplitude_at_1.5h, amplitude_at_0.58h, amplitude_at_8.83h | 1.02 |
| 5 | integral_at_19.08h, amplitude_at_1.5h, amplitude_at_0.58h, amplitude_at_8.83h, envelope_above_2.06 | 1.05 |
| 6 | integral_at_19.08h, amplitude_at_1.5h, amplitude_at_0.58h, amplitude_at_8.83h, envelope_above_2.06, duration_above_2.27 | 1.05 |
| 7 | integral_at_6.75h, amplitude_at_1.5h, amplitude_at_11h, 1st_peak_amplitude, amplitude_at_0h, amplitude_at_8.83h, amplitude_at_0.58h | 1.06 |

**Supplementary Table 4E: Channel capacity measurements & metrics in optimized vectors across doses of LPS**

| Dim | Metrics | Channel Capacity |
|---|---|---|
| 1 | integral_at_2.25h | 0.91 |
| 2 | integral_at_2.25h, amplitude_at_0.33h | 1.20 |
| 3 | integral_at_2.25h, amplitude_at_0.33h, amplitude_at_0.5h | 1.30 |
| 4 | integral_at_2.25h, amplitude_at_0.33h, amplitude_at_0.5h, duration_above_1.65 | 1.37 |
| 5 | integral_at_2.25h, amplitude_at_0.33h, amplitude_at_0.5h, duration_above_1.65, amplitude_at_0.25h | 1.40 |
| 6 | integral_at_2.25h, amplitude_at_0.33h, amplitude_at_0.5h, duration_above_1.65, amplitude_at_0.25h, amplitude_at_11.5h | 1.44 |
| 7 | integral_at_2.25h, amplitude_at_0.33h, amplitude_at_0.5h, duration_above_1.65, amplitude_at_0.25h, amplitude_at_11.5h, amplitude_at_0h | 1.45 |

**Supplementary Table 4F: Channel capacity measurements & metrics in optimized vectors across doses of CpG**

| Dim | Metrics | Channel Capacity |
|---|---|---|
| 1 | integral_at_2.08h | 0.89 |
| 2 | integral_at_2.75h, amplitude_at_0.25h | 0.95 |
| 3 | integral_at_2.75h, amplitude_at_0.25h, integral_at_1.75h | 0.99 |
| 4 | integral_at_2.75h, amplitude_at_0.25h, integral_at_1.75h, envelope_above_2.06 | 1.00 |
| 5 | integral_at_2.75h, amplitude_at_0.25h, integral_at_1.75h, envelope_above_2.06, amplitude_at_10.17h | 1.01 |
| 6 | integral_at_2.75h, amplitude_at_0.25h, integral_at_1.75h, envelope_above_2.06, amplitude_at_10.17h, amplitude_at_0.08h | 1.02 |
| 7 | integral_at_2.75h, amplitude_at_0.25h, integral_at_1.75h, envelope_above_ 2.06, amplitude_at_10.17h, amplitude_at_0.08h, envelope_above_0.82 | 1.03 |

**Supplementary Note 1.**
Details for channel capacity calculations and codeword Identification

We used Shannon's information theoretic framework to correlate the stimulus condition to dynamical features extracted from temporal trajectories of NFκB activity.

$$\begin{array}{c} noise \\ \downarrow \\ \boldsymbol{X} \rightarrow \boxed{communication\ channel} \rightarrow \boldsymbol{Y} \end{array}$$

$\boldsymbol{X} = stimulus\ condition$
$\boldsymbol{Y} = NF\kappa B\ dynamical\ features$
$C(Q) = \max_{P_x} I(\boldsymbol{Y};\boldsymbol{X})$

$$I(\boldsymbol{Y};\boldsymbol{X}) = H_{diff}(\boldsymbol{Y}) - H_{diff}(\boldsymbol{Y}|\boldsymbol{X})$$
$$H_{diff}(\boldsymbol{Y}|\boldsymbol{X}) = \Sigma_{i=1}^{m} q_i H_{diff(Y_i|X=x_i)} = -\Sigma_{i=1}^{m} q_i \Sigma_{j=1}^{n_i} \frac{1}{n_i} log2(f(Y_i = y_{ij}|X = x_i))$$
$$H_{diff}(Y) = -\Sigma_{i=1}^{m} \frac{q_i}{n_i} \Sigma_{j=1}^{n_i} \log_2(f\left(\boldsymbol{Y} = y_{i_j}\right))$$
$$f(\boldsymbol{Y} = y) = \Sigma_{w=1}^{m} q_w f(\boldsymbol{Y} = y \mid \boldsymbol{X} = x_w)$$

$$H_{diff}(\boldsymbol{A}) = -\Sigma_{j=1}^{N_a} \delta_j \log_2 f(a_j)),\ where\ \delta_j = probability\ of\ observating\ a$$
$$f(a_j|\boldsymbol{A}) = \frac{k}{N_a V_d z(a_j|\boldsymbol{A}))^{\frac{d}{k}}}$$
$$V_d = \frac{\pi^{\frac{d}{2}}}{\Gamma\left(\frac{d}{2}+1\right)}$$

$H_{diff}(\boldsymbol{Y}|\boldsymbol{X}) = conditional\ entropy$
$m = number\ of\ stimulus\ conditions$
$n = number\ of\ cells\ in\ a\ condition$
$q_i = probability\ of\ observing\ a\ stimulus$
$x_{ij} = a\ single\ cell's\ response$
$k = number\ of\ neighbors\ used\ in\ kNN\ estimate\ of\ marginal\ distribution\ of\ Y$
$d = vector\ dimension$
$\delta_j = probability\ of\ observation$

*Controlling for different sample sizes.* Jackknife resampling was used to control for different sample sizes by calculating channel capacity for differently-sized subsets and extrapolating to an infinite sample size.

$n_c = 24$
*Setting threshold.*

1. $\mathbf{t} \leftarrow \left(\frac{1}{\sqrt{2}}\right)^{[1:6]} - \left(\frac{1}{2}\right)^{[1:6]}$
2. $t_1 \leftarrow 0.3$
3. If $d > 6$ then $\mathbf{t} \leftarrow [\mathbf{t}, 0.1 * \mathbf{1_{d-6}}]$

$$For\ i = 1 \ldots d$$

1. Compute channel capacity by optimizing over marginal distribution of **X**
   a. For $j$ = 1… $k$
      i. $c_j \leftarrow \max_{p_X} I(\boldsymbol{x_j}; \boldsymbol{Y})$
      ii. $\boldsymbol{q_j} \leftarrow argmax_{PX} I(\boldsymbol{x_j}; \boldsymbol{Y})$

2. Select a subset of feature vectors whose channel capacity values exceeds $t_i$
   a. $\boldsymbol{X}^* \leftarrow \{\boldsymbol{x_j} \mid c_j > t_i\}$
   b. $\boldsymbol{Q}^* \coloneqq argmax_{PX} I(\boldsymbol{X}^*; \boldsymbol{Y})$

3. Select a subset of feature vectors that maximizes diversity of marginal distributions
   a. Select feature vector that yields the maximum channel capacity
      $\widehat{\boldsymbol{x}} \leftarrow \{\boldsymbol{x_j^*} \mid c_j = \max(\mathbf{c})\}$ , equivalently $\widehat{\boldsymbol{x}} \leftarrow argmax_{\boldsymbol{X}^*} \max_{p_X} I(\boldsymbol{X}; \boldsymbol{Y})$
   b. Construct a set of feature vectors containing the $\widehat{\boldsymbol{x}}$ and feature vectors whose marginal distributions, $\boldsymbol{q_j}$, are most orthogonal to $\widehat{\boldsymbol{q}} \leftarrow argmax_{PX} I(\widehat{\boldsymbol{x}}; \boldsymbol{Y})$
   c. $\boldsymbol{x_1^o} \leftarrow \widehat{\boldsymbol{x}}$ , $\boldsymbol{q_1^o} \leftarrow \widehat{\boldsymbol{q}}$
      i. $For\ m = 2 \ldots n_c$
         1. $\boldsymbol{Q^c} \coloneqq \{\boldsymbol{q} | \boldsymbol{q} \in \boldsymbol{Q}^* \wedge \boldsymbol{q} \notin \boldsymbol{Q}^o\}$
         2. $\boldsymbol{q_m^o} \leftarrow argmin_{\boldsymbol{Q}^c} \|\boldsymbol{Q^o} - \boldsymbol{Q^c}\|_2$
         3. $\boldsymbol{x_m^o} \leftarrow \{\boldsymbol{x_j^*} | \boldsymbol{q_j^*} \equiv \boldsymbol{q_m^o}\}$
